# Large-scale template-based structural modeling of T-cell receptors with known antigen specificity reveals complementarity features

**DOI:** 10.1101/2023.03.29.533758

**Authors:** Dmitrii S. Shcherbinin, Vadim K. Karnaukhov, Ivan V. Zvyagin, Dmitriy M. Chudakov, Mikhail Shugay

## Abstract

T-cell receptor (TCR) recognition of foreign peptides presented by the major histocompatibility complex (MHC) initiates the adaptive immune response against pathogens. A large number of TCR sequences specific to different antigens are known to date, however, the structural data describing the conformation and contacting residues for TCR:antigen:MHC complexes is relatively limited. In the present study we aim to extend and analyze the set of available structures by performing highly accurate template-based modeling of TCR:antigen:MHC complexes using TCR sequences with known specificity. Using the set of 29 complex templates (including a template with SARS-CoV-2 antigen) and 732 specificity records, we built a database of 1585 model structures carrying substitutions in either TCRα or TCRβ chains with some models representing the result of different mutation pathways for the same final structure. This database allowed us to analyze features of amino acid contacts in TCR:antigen interfaces that govern antigen recognition preferences and interpret these interactions in terms of physicochemical properties of interacting residues. Our results provide a methodology for creating high-quality TCR:antigen:MHC models for antigens of interest that can be utilized to predict TCR specificity.

## Introduction

Specific interaction between T-cell receptors (TCRs) and major histocompatibility complex (MHC)-peptide complexes is a key initiation factor of adaptive immunity-driven physiological processes (*1*) that happens upon recognition of an infected or antigen-presenting cell by cognate T-cell. TCR molecules are heterodimers encoded by genes formed via the process of V(D)J-rearrangement that ensures the presence of a highly diverse (>10^8^ combinations of TCRα - TCRβ chain pairs) TCR sequences in an individual, required to recognize and target previously unencountered pathogens viewed through the lens of MHC molecules presenting peptides (*2,3*). Describing molecular mechanisms governing antigen recognition is therefore the cornerstone of adaptive immunity studies making it possible to predict immune responses to specific antigens, as well as cross-reactivity and selectivity of TCRs in the near future (*4*). Moreover, present methods, even high-throughput ones such as 10X single-cell sequencing with dCODE dextramers (5), cannot yield enough TCR specificity data to cover the space of all possible TCR:pMHC interactions.

Currently, several databases gathering experimental information about TCR-peptide recognition are available to the research community: VDJdb, IEDB, McPAS-TCR (6–8). These resources store and aggregate data on primary TCR sequences, cognate epitopes and MHC context mined using literature search.

In addition there are more than 280 crystal structures stored in the Protein Data Bank (PDB) that encode the geometry and physical contacts in TCR-pMHC complexes (9,10). Several computational approaches have been developed for homology-based modeling of individual TCR structures or TCRs in their ternary complex with peptide-MHC molecules they recognize. Promising software tools and approaches in this area include the TCRmodel (11) and TCR-pMHC-models (12) services.

In addition to homology modeling approaches, it has been shown that AlphaFold software, which is based on deep learning techniques and has revolutionized the field of protein structure prediction, can be used for the *de-novo* modeling of TCR complexes with pMHC, although the proposed pipeline needs further tuning, especially for CDR3 loops (13).

Recent studies, focused on structural bioinformatics and molecular modeling of TCR-epitope-MHC complexes, can be categorized into two primary areas of research. The first area involves large-scale modeling approaches and mathematical modeling of a significant amount of structural data, while the second area is devoted to a detailed examination of individual complexes and enhancing the precision of binding specificity and affinity inference.

Large-scale attempts to model multiple TCR:pMHC complexes have been carried out in a number of recent studies, most of them were focused on serving as a proof-of-concept and as a benchmark for modeling software, not directly aimed at general applications, such as studying regularities in TCR:pMHC recognition at the residue level or imputing a missing TCR:pMHC structure for a TCR linked to a certain disease. A consistent database of over 23,000 TCR complex structures with pMHC was created using template-based approaches for TCR (Repertoire Builder (14), LYRA (15)), MHC (NetMHCpan (16)) modeling, and construction of docked complexes (17).

Machine learning algorithms have demonstrated their usefulness in predicting TCR-peptide interactions employing high similarity (HS) and random sampling strategies for deep learning models (18) and evaluating binding affinity by training a random forest classifier model using the ATLAS database (19).

On the other hand, it was demonstrated using two contrasting TCR−pHLA test sets that the MMPB/GBSA approach can be effectively applied during the binding affinity calculation.However, the protocol needs to be adjusted based on the similarity of the compared structures (20). Analysis of multiple molecular docking approaches (21) and their corresponding scoring functions has indicated that hydrophobic and electrostatic interactions play important roles in TCR-pMHC recognition. Additionally, a priori knowledge of contacting residues of TCR and peptide has been shown to improve modeling of the entire complex, including highly flexible CDR3 loops.

The importance of hydrophobic interactions and pi-stacking forces in TCR contacts with HLA-A*02-restricted peptides, was also analyzed and illustrated using molecular dynamics simulations and binding free energy calculations (22).

Despite the fact that much attention has been paid to the study of SARS-CoV2 recently, only a few COVID-associated TCRs have a resolved TCR:pMHC structure to date (23–26). Having a variety of structures with SARS-CoV-2 epitopes can aid in studying T-cell recognition of infected cells, especially since the number of known COVID-associated TCRs is constantly growing (6).The study of TCR – peptide MHC complexes is important in understanding the immune response to SARS-CoV-2 vaccinations and the viral immune evasion strategies employed by VOCs (Variants of concern). Mutations in T cell epitopes can disrupt recognition by TCRs, which highlights the importance of modeling and studying the spatial structures of TCR – peptide MHC complexes.

There are several main factors that guide antigen recognition in TCR:pMHC complex: placement of the peptide in the MHC groove, pairing of TCR-alpha and beta chains, specific orientation of pMHC complex against T-cell receptor and selective interaction of TCR residues with MHC and peptide.

Most studies suggest that TCRs recognize peptide-MHC complexes via CDR1-3 loops. While CDR1 and CDR2 loops stabilize binding with MHC molecules and target small parts of a protein, most of the interaction between TCR and antigen occurs via CDR3, which are omega-loops of 10-20 amino acids (27,28). Also, it has been shown that CDR3 loop sequence’s mid-region creates most of the contacts with peptide due to the loop and peptide geometry. Additionally, both alpha and beta chains contribute to antigen recognition, although sometimes this contribution is disproportionate (29).

The present study reports the results of large-scale template-based modeling of the VDJdb, which is a curated database of TCR sequences with known specificities. We aim at traversing the set of TCR:pMHC specificity records that differ by a fixed number of amino acid substitutions from the original template, as it was shown that 1-3 consequent substitutions rarely impact antigen specificity (30). Our primary aim is to establish a pipeline that can be easily employed to create accurate and reliable spatial structures for studies that report massive sequencing of TCRs specific to a panel of antigens. By applying this pipeline to the existing database we aim at providing access to structural data for immunologists and biologists unfamiliar with *in silico* structural modeling. We also demonstrate, as a proof-of-concept, that such modeling can be used to study features of residue interactions in TCR:pMHC complexes, and report statistics of various physicochemical interactions between TCR and peptide.

## Results

### Building a compendium of TCR:pMHC models for VDJdb

In this study we developed a bioinformatic pipeline for modeling T cell receptor CDR3 loops by stepwise introduction of single amino acid mutations to known spatial structures of complexes used it to construct 1585 TCR-peptide-MHC models and analyzed contacting residues between CDR3 loops and peptides in modeled complexes. The pipeline applied in our approach is shown in **Figure 1 (A** and **B)**. In the first step we curated a pre-processed and annotated (see Methods) database of TCR-pMHC structures from PDB and built clusters of annotated CDR3 sequences from VDJdb, using CDR3 loops from known PDBs as cores. In the next stage we calculated pathways for consequent introduction of single amino acid substitutions to cover all sequences in selected clusters, starting from known PDB structures. The actual extent of VDJdb coverage by our modeling approach is depicted in **Figure 1C**, and the modeling paths are presented in **Supplementary Figure S1**.

**Figure 1.**
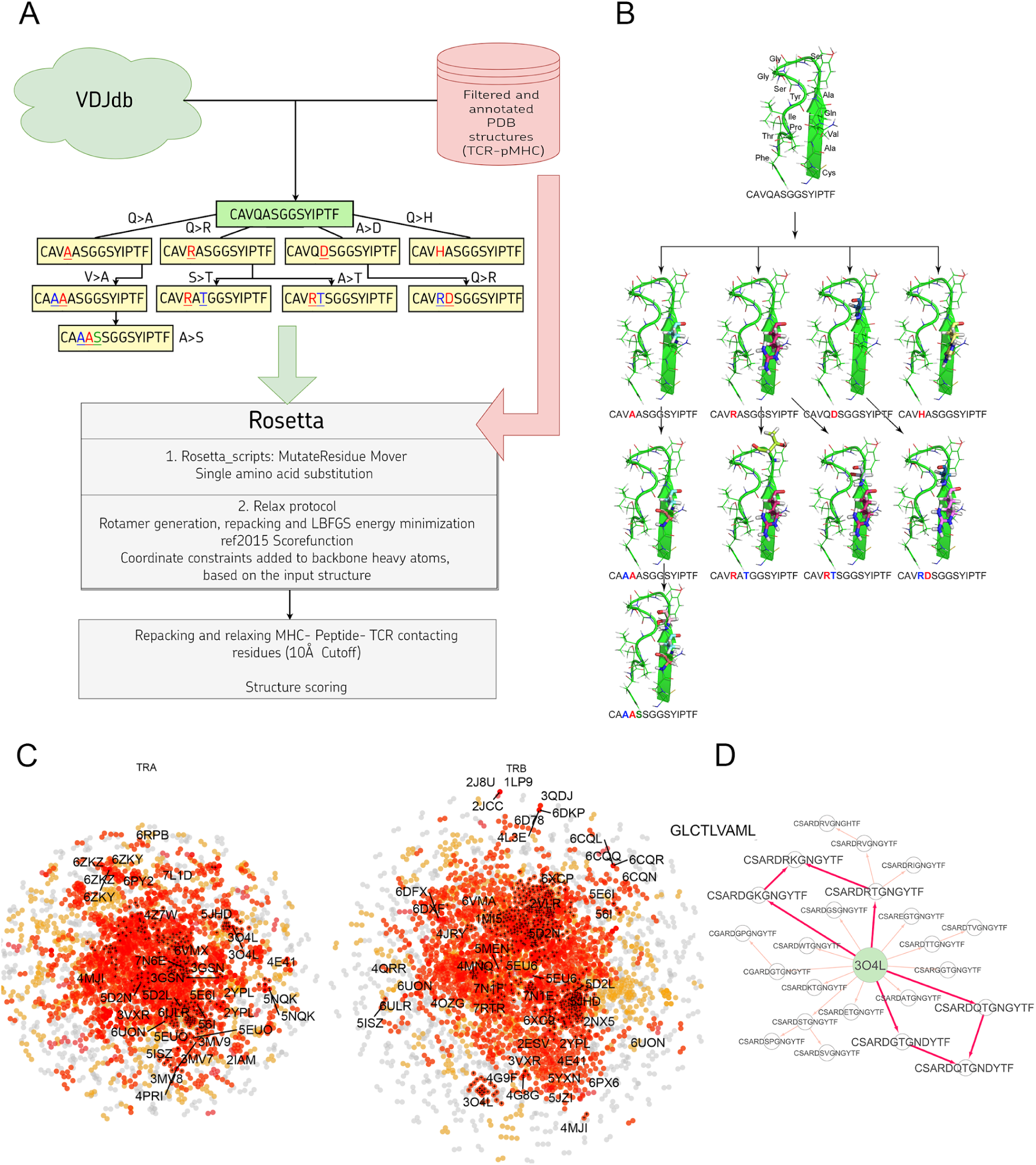
A) Pipeline used in the stepwise amino acid mutation modeling approach applied in the present study. B) Mutation path in one of the studied CDR3 clusters starting from a known 2NX5 PDB structure (31) carrying CAVQASGGSYIPTF CDR3α loop. C) CDR3 sequence similarity map of VDJdb, layout of a graph with edges connecting CDR3 sequences that differ by no more than a single amino acid mismatch. CDR3 sequences having PDB templates are shown with labels. Orange points show VDJdb entries that were classified as belonging to a sequence homology motif (described in (32)), red points are connected components built around PDB structure templates, TCR:pMHC records connected by no more than 3 subsequent amino acid substitutions to a PDB that were modeled in present study are shown with black crosses. D) An example of the mutation pathways in which the same CDR3 loops could be modeled using different intermediate sequences. The corresponding pathways are shown with bold red arrows.

Using 1030 unique mutation pathways, all initial structures were altered and further optimized using Rosetta protocols. As a result, we built 1585 models based on 29 template structures from the PDB database, which contain 247 unique sequences of CDR3 loops for TCR-alpha and 485 for TCR-beta chains. The total number of modeled structures is greater than the number of unique pathways or unique CDR3 sequences because the same CDR3 loops carrying, for example, two or three substitutions can be modeled through various intermediate sequences. Additionally, different initial PDB templates could be used to model the same CDR3 loops. All modeled CDR3 sequences are presented in **Supplementary Figure S2**, and the number of models according to TCR chains, presence of contact with peptide, and type of contact are presented in **Supplementary Table S1**.

The initial templates are presented in **Table1**. It is worth noting that some PDB templates carried identical TCR chains. During further analysis, models built based on such templates were grouped, and their calculated spatial and energy properties were averaged. These grouped structures would be referred to as “non-redundant”.

### Analysis of CDR3:peptide contacts in modeled structures

Using a 5Å cutoff distance, contacting (or “interacting”) residues of “mutated” CDR3 loops and peptides were identified in all modeled structures and structures obtained from the PDB database. It was shown that only about half of them carried substitution in contacting positions: approximately 48% (763 models out of 1585) in all TCR-pMHC complex models and ∼51% (551 models out of 1085) in the non-redundant set of structures.

Since all the modeled TCRs in the cluster have the same specificity as TCR in the core crystal structure, we expect that modeled amino acid substitutions should not dramatically influence the binding between TCR and epitope. That might be the reason why many amino acid variations were found in non-contacting positions - they are simply distal and do not alter TCR-pMHC interface. Additionally, it can be assumed that the remaining substitutions in contacting positions should not be critical for the affinity level of binding between CDR3 loops and epitopes.

CDR3–epitope interactions were analyzed in all initial templates, and for each structure the most valuable positions in CDR3 loops were identified according to the most significant energetic impact (calculated residue-wise energy values) of corresponding amino acids on TCR-epitope binding. These results were compared with estimated frequencies of amino acid variations in each position on CDR3 loops in our modeled structures. It was found out that only 11% of all mutations and 23% of mutations in contacting residues were in energetically valuable positions. In the non-redundant set the rate was slightly higher: 15% and 29% respectively. These results also indicate that substitutions mainly occur in regions of CDR3 loops that are not very important for epitope recognition, thus preserving specificity and affinity of binding in TCR-pMHC complexes.

All amino acid pairs involved in CDR3-peptide interactions were identified and counted in all modeled and 152 original structures of human TCR-peptide-MHCI complexes. Interactions between amino acid residues of CDR3 loops and peptides were classified based on their distances and contacting parts: main chain or side chain groups. We considered either all contacts (a single TCR residue can contact with several peptide residues located in distance <5Å) or only closest contacting residues (for each TCR residue only a single contact with the closest residue in the peptide was considered). Four groups were formed based on these parameters: (1) all contacting residue pairs, (2) closest contacting residues (for each TCR residue only a single contact with the closest residue in the peptide was considered), (3) residues contacting through side chain groups, and (4) closest contacting residues through side chain groups. For the third and fourth groups we considered only contacts in which at least one of the residues interact through a side chain group. Analysis of side chain groups is crucial because practically all physicochemical features of amino acid residues are “encoded” in side chain groups.

The number of all found contacting amino acid pairs between CDR3 loops and peptides in original and modeled structures is presented in **Figure 2**.

**Figure 2.**
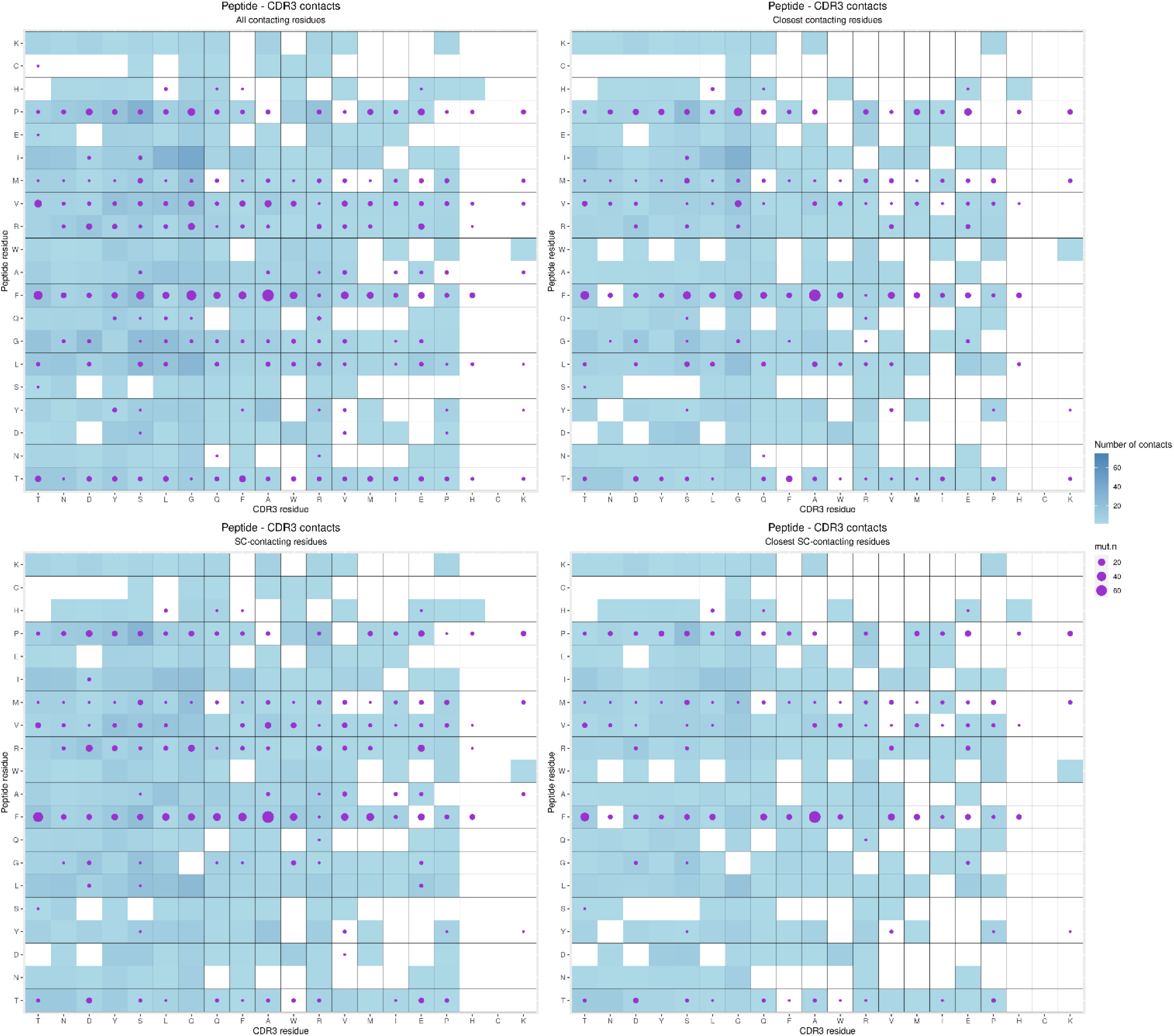
Contacting amino acid pairs identified in original and modeled structures. Contacts in the original structures are represented as squares, while contacts in the modeled structures are represented as circles. The number of identified contacting pairs is illustrated as a gradient fill for the original structures, and the size of circles represents the results for the modeled structures. The results for all contacting residue pairs are presented in the upper-left corner; closest contacting residue pairs are shown in the upper right corner. Amino acid residues, contacted by at least one side chain group are in the down-left corner, and the results for closest amino acid residues contacting by at least one side chain group are presented in the down-right corner.

It can be seen that, even though there are some new contacting residue pairs in the modeled structures that were not found in the original structures, the majority was not unique.

We visually inspected the contacting residues in the CDR3 alpha and beta clusters modeled from the core sequences presented in 7RTR and 7N1F original PDB structures. These structures contain TRAV12-2*01 / TRAJ30*01 TCRα chain with “CAVNRDDKIIF” CDR3, TRBV7-9*01 / TRBJ2-7*01 TCRβ chain with “CASSPDIEQYF” CDR3, and YLQPRTFLL SARS-CoV2 epitope presented by MHCI. Using selected clusters, 54 unique single amino acid substitutions were modeled: 16 substitutions were introduced to the CDR3 loop of TCR alpha and 38 to TCR beta.

Analyzing all these modeled substitutions, it was found out that in most cases they were in non-contacting positions according to the corresponding original structures. Contacting residues were only changed to similar amino acids, while more dissimilar ones only appeared in positions where contact with the peptide was formed by main chain atoms. Therefore, changes of side chain groups should not influence CDR3-peptide binding.

The modeled substitutions are shown in **Figure 3**, and corresponding information about the contacting residues between the CDR3 loops and peptide is presented in **Table 2**.

**Figure 3.**
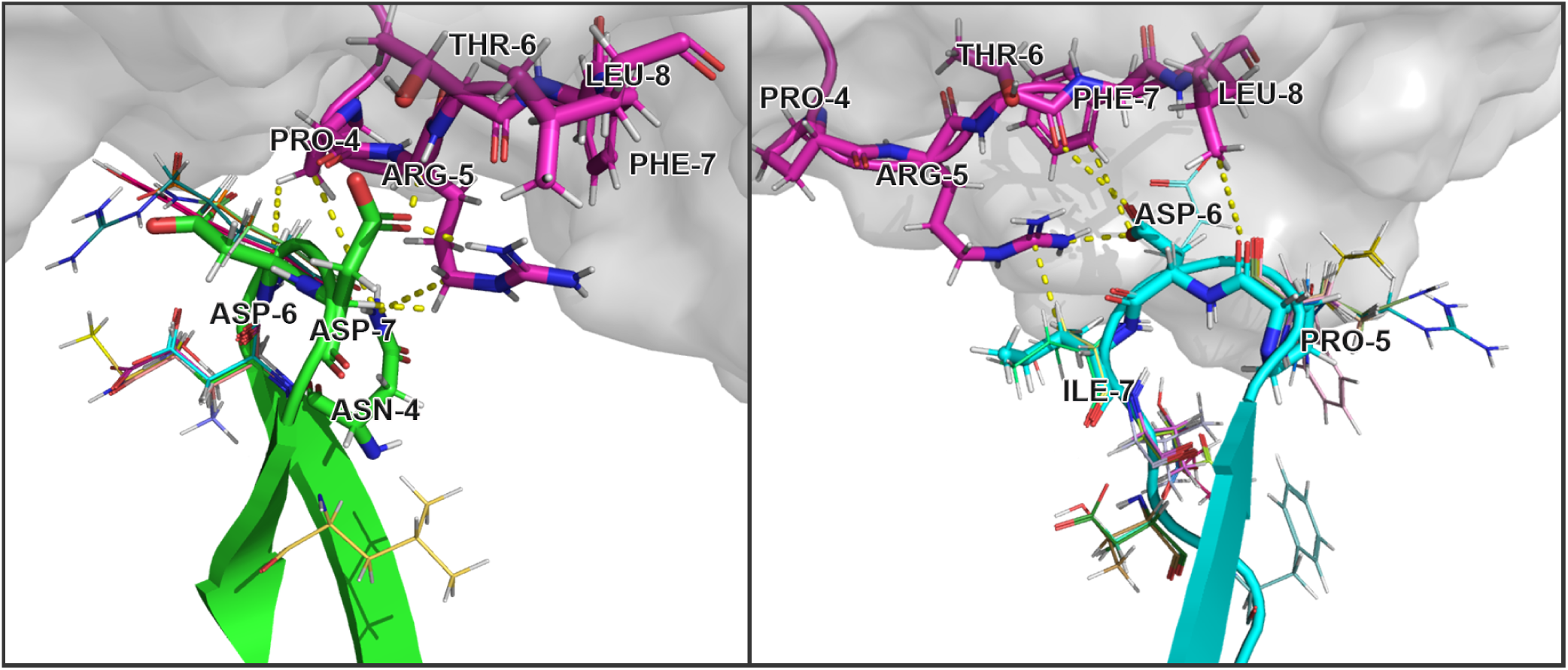
Modeled amino acid substitutions in CDR3 loops in TCR:peptide:MHC complexes containing SARS-CoV-2 spike epitopes YLQPRTFLL. Contacting residues are represented as sticks, modeled substitutions are represented as lines and close contacts are illustrated as dashed sticks. CDR3 loop of TCRα is colored green, TCRβ is colored cyan, and the peptide residues are colored magenta.

**Table 1.**
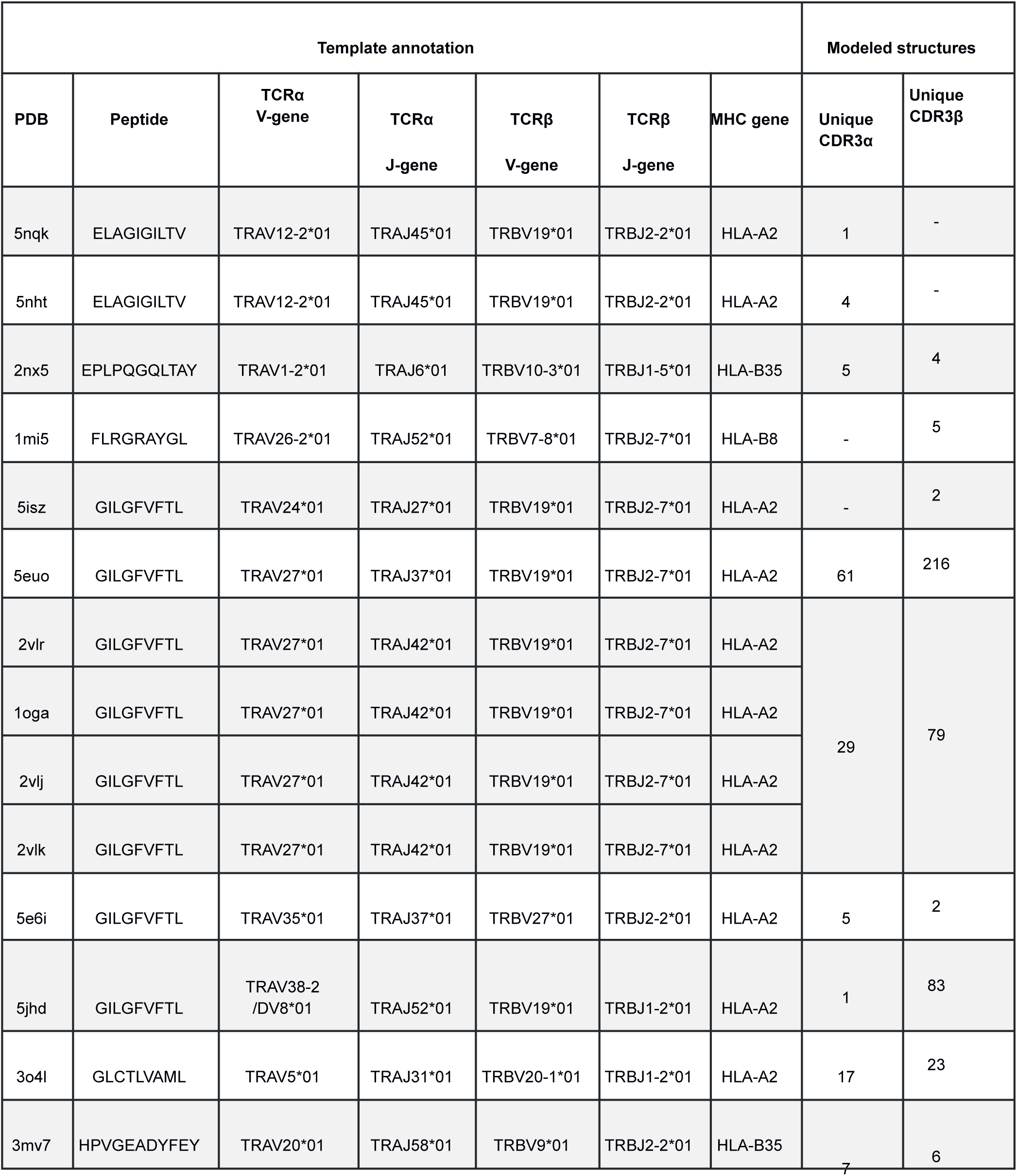
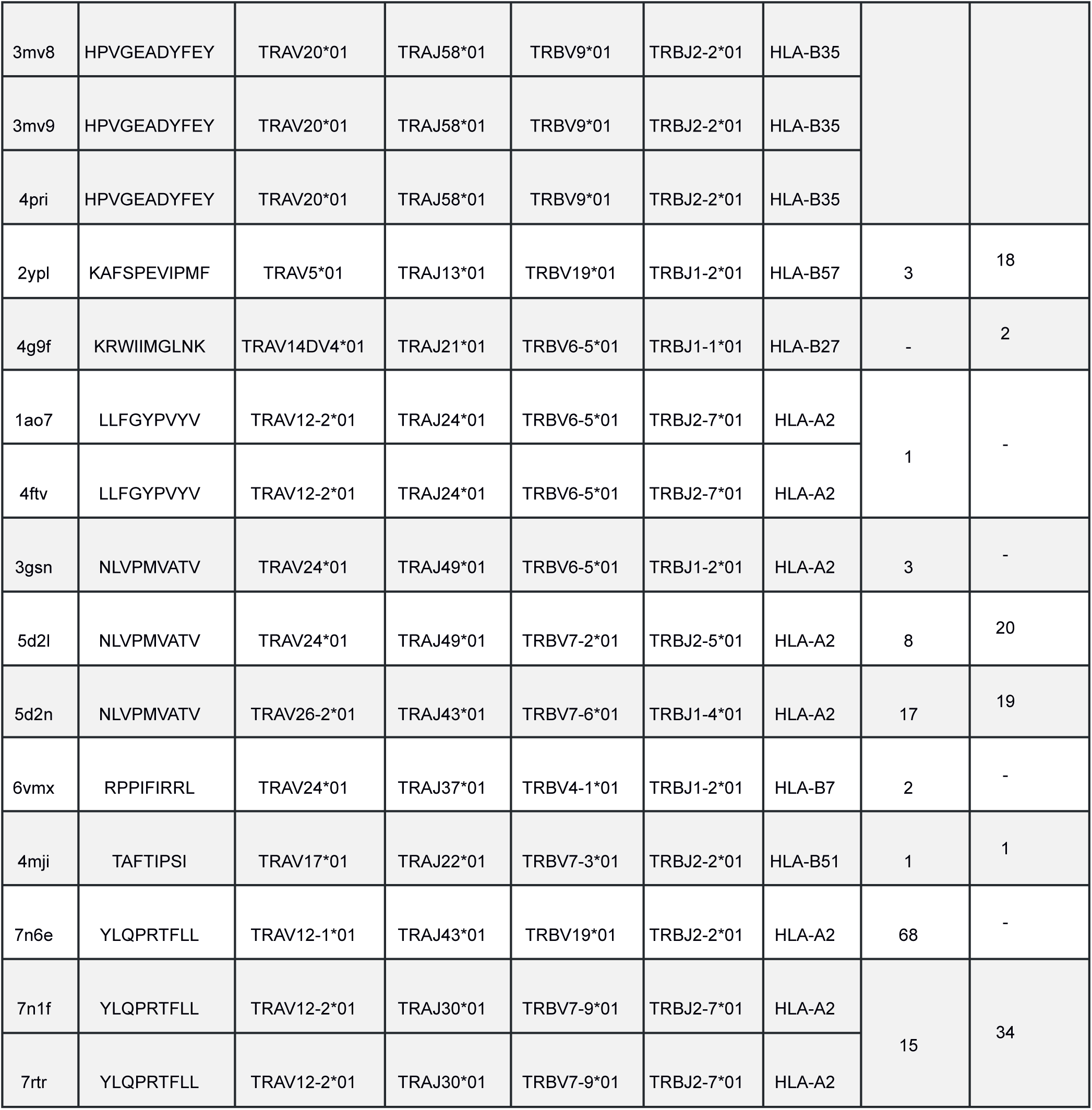
Details for initial PDB templates and the number VDJdb entries modeled as structures. *Dash (“-”) indicates there were no VDJdb records with 1-3 mismatches we could model using our approach*.

**Table 2.** Contacting residues in modeled TCR:peptide:MHC complexes, built based on 7RTR and 7N1F original PDB structures. *Annotation of interactions is presented in accordance with the results of the analysis of residue interaction networks. Provided amino acid numbering starts from the beginning of the corresponding substructure – CDR3 loop or peptide*.\

| CDR3 |  |  | Peptide |  | Interaction |  |  | Modeled substitutions |
| --- | --- | --- | --- | --- | --- | --- | --- | --- |
| chain | residue | position | residue | position | Type of interaction | cdr3 | peptide |  |
| TCRα | Asn | 4 | Pro | 4 | VDW | side chain | side chain | none |
|  |  |  | Arg | 5 | VDW | side chain | side chain |  |
|  | Asp | 6 | Pro | 4 | VDW | main chain | main chain | multiple |
|  |  |  | Arg | 5 | VDW | main chain | side chain |  |
|  | Asp | 7 | Pro | 4 | VDW | side chain | main chain | none |
|  |  |  | Arg | 5 | HBOND | side chain | side chain |  |
|  |  |  | Thr | 6 | HBOND | side chain | side chain |  |
| TCRβ | Pro | 5 | Leu | 8 | VDW | main chain | side chain | multiple |
|  | Asp | 6 | Arg | 5 | IONIC | side chain | side chain | GLU |
|  |  |  | Thr | 6 | VDW | side chain | main chain |  |
|  |  |  | Phe | 7 | VDW | side chain | side chain |  |
|  | Ile | 7 | Arg | 5 | VDW | side chain | side chain | VAL, SER |

During the inspection of the selected modeled structures (**Table 2**) it was observed that only two contacting positions carried multiple variations of amino acids: Asp-6 in the CDR3α and Pro-5 in the CDR3β loop. In both cases, the side chain groups of the mutated residues were oriented towards the MHC molecule instead of the epitope. Two other positions carried one or two variants of substituted amino acids that were quite similar to the original in terms of their physicochemical properties: Glu instead of Asp in the 6th position and Val or Ser instead of ILE in the 7th position of CDR3β loop.

### Correlation of CDR3:peptide binding energy values and amino acid properties

It was previously shown (33) that the most important energy terms during the calculation of free binding energies between TCRs and peptides were the attractive van der Waals impact energy, solvation energy, and side-chain – side-chain hydrogen bond energy, while the repulsive van der Waals term was insignificant. Thus, in our research, energy values for contacting residues and interfaces between modified CDR3 loops and peptides in all modeled complexes were calculated using full Rosetta scoring functions and two of its limited variations: “Large patch” and “Small patch” presets (see the “Methods” section). As all models were built using stepwise introduction of single amino acid substitutions, we calculated delta values of free energies of interacting interfaces between complexes differing by 1 amino acid. These energy values represent the impacts of point amino acid variations on the affinity of complexes. Henceforth that type of values would be referred to as “dEnergy”.

#### Correlation between BLOSUM scores and dEnergy values

Two types of BLOSUM matrices (BLOSUM62 and BLOSUM100) were used to study the consistency between amino acid substitution scores and their impact on dEnergy in modeled complexes. Pearson’s correlations were calculated between absolute values of free energy adjustments caused by mutation and one of three modified BLOSUM indices of corresponding substitutions: Clustered Target Frequencies (QIJ), Clustered Scoring Matrix in Bit Units (SIJ) or most commonly used Clustered Scoring Matrix in 1/2 Bit Units (BLA). Adjusted index values were calculated as follows:

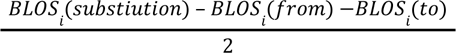

Where *BLOS_i_*(substitution)- BLOSUM index of studied substitution

*BLOS_i_*(from) - BLOSUM index of a match of origin residue

*BLOS_i_*(to)- BLOSUM index of a match of new residue

As BLOSUM scores represent the ratio of the likelihood of two amino acids being exchanged with biological significance, it is more reasonable to compare them with absolute values of dEnergy without taking into consideration its sign. Thus, we can evaluate the impact of substitutions to similar or dissimilar amino acids. Energy values were calculated using the full Rosetta scoring function and “Large patch” and “Small patch” presets.

Correlations were analyzed for all mutated residues in CDR3 loops and separately for substitutions in only contacting positions. Top 20 corresponding correlation values and p-values are presented in **Table 3**.

**Table 3.** Correlation values, calculated for BLOSUM scores and absolute values of dEnergy after the introduction of amino acid substitutions in modeled complexes.

|  | All mutations |  | Contacting mutations |  |  |  |
| --- | --- | --- | --- | --- | --- | --- |
| Blosum index | Pearson correlation | P-value | Pearson correlation | P-value | Energy scoring | Optimization |
| BLA.62.v2 | -0.136 | **2.03E-04** | -0.219 | **7.76E-06** | Small patch | Minimized |
| BLA.62.v2 | -0.135 | **2.36E-04** | -0.205 | **2.69E-05** | Large patch | Minimized |
| SIJ.62.v2 | -0.112 | **0.002** | -0.182 | **2.16E-04** | Small patch | Minimized |
| SIJ.100.v2 | -0.097 | **0.008** | -0.165 | **6.67E-11** | Small patch | Minimized |
| SIJ.62.v2 | -0.107 | **0.004** | -0.164 | **0.001** | Large patch | Minimized |
| BLA.100.v2 | -0.096 | **0.009** | -0.164 | **1.86E-10** | Small patch | Minimized |
| BLA.62.v2 | -0.135 | **4.34E-05** | -0.161 | **3.07E-04** | Large patch | Repacked |
| SIJ.100.v2 | -0.084 | **0.022** | -0.137 | **0.005** | Large patch | Minimized |
| BLA.100.v2 | -0.083 | **0.024** | -0.136 | **0.006** | Large patch | Minimized |
| BLA.62.v2 | -0.123 | **1.79E-04** | -0.133 | **0.003** | Small patch | Repacked |
| BLA.62.v2 | -0.055 | 0.119 | -0.128 | **0.008** | Rosetta | Minimized |
| SIJ.62.v2 | -0.117 | **4.01E-04** | -0.126 | **0.005** | Large patch | Repacked |
| SIJ.62.v2 | -0.050 | 0.154 | -0.113 | **0.019** | Rosetta | Minimized |
| SIJ.100.v2 | -0.107 | **0.001** | -0.110 | **0.014** | Large patch | Repacked |
| SIJ.62.v2 | -0.082 | **0.011** | -0.109 | **0.015** | Rosetta | Repacked |
| SIJ.100.v2 | -0.085 | **0.009** | -0.102 | **0.023** | Rosetta | Repacked |
| BLA.100.v2 | -0.106 | **0.001** | -0.099 | **0.026** | Large patch | Repacked |
| BLA.100.v2 | -0.038 | 0.287 | -0.099 | **0.040** | Rosetta | Minimized |
| BLA.62.v2 | -0.071 | **0.028** | -0.097 | **0.029** | Rosetta | Repacked |
| SIJ.100.v2 | -0.040 | 0.261 | -0.097 | **0.044** | Rosetta | Minimized |

It was observed that the correlations between different BLOSUM62 or BLOSUM100 indexes and absolute delta energy values were very poor overall. The best correlation was observed between BLA.62 values and the absolute value of dEnergy, calculated using the “Small patch” preset in structures that carried mutations in contacting positions. Pearson correlation values (R coefficient) was −0.22 and −0.13 for minimized and repacked structures respectively. The results are presented in **Fig. 4**. It can also be observed that correlation is better for TCRβ chains, which corresponds to previously shown results that beta-chains are more crucial for the specificity of peptide recognition and binding.

**Figure. 4.**
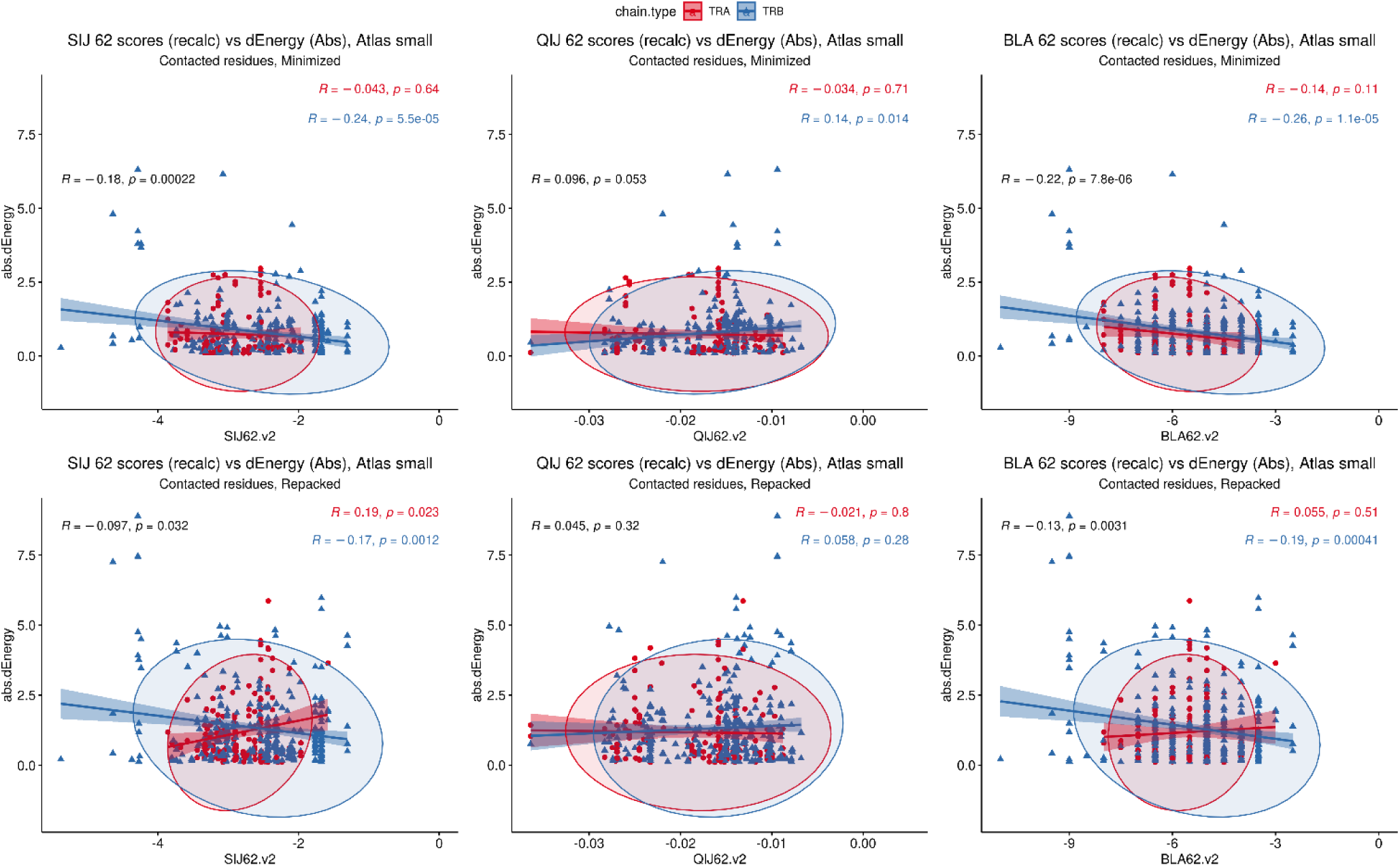
Correlation of BLOSUM62 matrix indices and absolute values of dEnergy, calculated for modeled structures. Corresponding values for TCRa and TCRb are colored red and blue.

These results show that standard BLOSUM matrices might not be as informative in the analysis of full CDR3 sequences, their clusterization and prediction of binding with specific peptides. The highest correlation was observed when only containing residues were considered. This fact highlights that instead of aligning and analyzing full CDR3 sequences, special attention should be paid to contacting parts.

Despite the low correlations overall, it was mentioned that using energy minimization as the final optimization of models led to a slight improvement of corresponding values compared to repacking. This can be explained by the “fitting” of contacting interfaces in TCR-pMHC complexes after repacking of “mutated” models, which resulted in “smoothing” (reduction) of the substitution impact.

#### Influence of remoteness of contacting amino acids on dEnergy

We additionally analyzed the impact of the distance between substituted residues and peptides on dEnergy of TCR-pMHC binding. In cases when mutated residue in CDR3 interacted with more than one residue of epitope, we used minimal distance values. It was shown that the total Pearson correlation values between distance and absolute values of dEnergy in minimized modeled structures were −0.27, −0.39 and −0.41 for energy values calculated using the full Rosetta scoring function, “Large-“ and “Small patch” presets respectively. The negativity in correlation values indicates that the more distant a modeled substitution is from the peptide, the less it impacts dEnergy values, which makes biophysical sense.

Analyzing these dependencies for TCRα and TCRβ separately, it can be seen that correlation was better for TCRβ in comparison with TCRα. These results indicate that CDR3 loops of beta-chains might have a greater impact on peptide recognition than TCR-alpha’s. Corresponding correlations, calculated for repacked structures were smaller: −0.074, −0.26 and −0.18. This decrease in correlation also supports the concept of “fitting” of mutated CDR3 residues after repacking optimization. Correlation graphs are presented in **Figure 5**.

**Figure 5.**
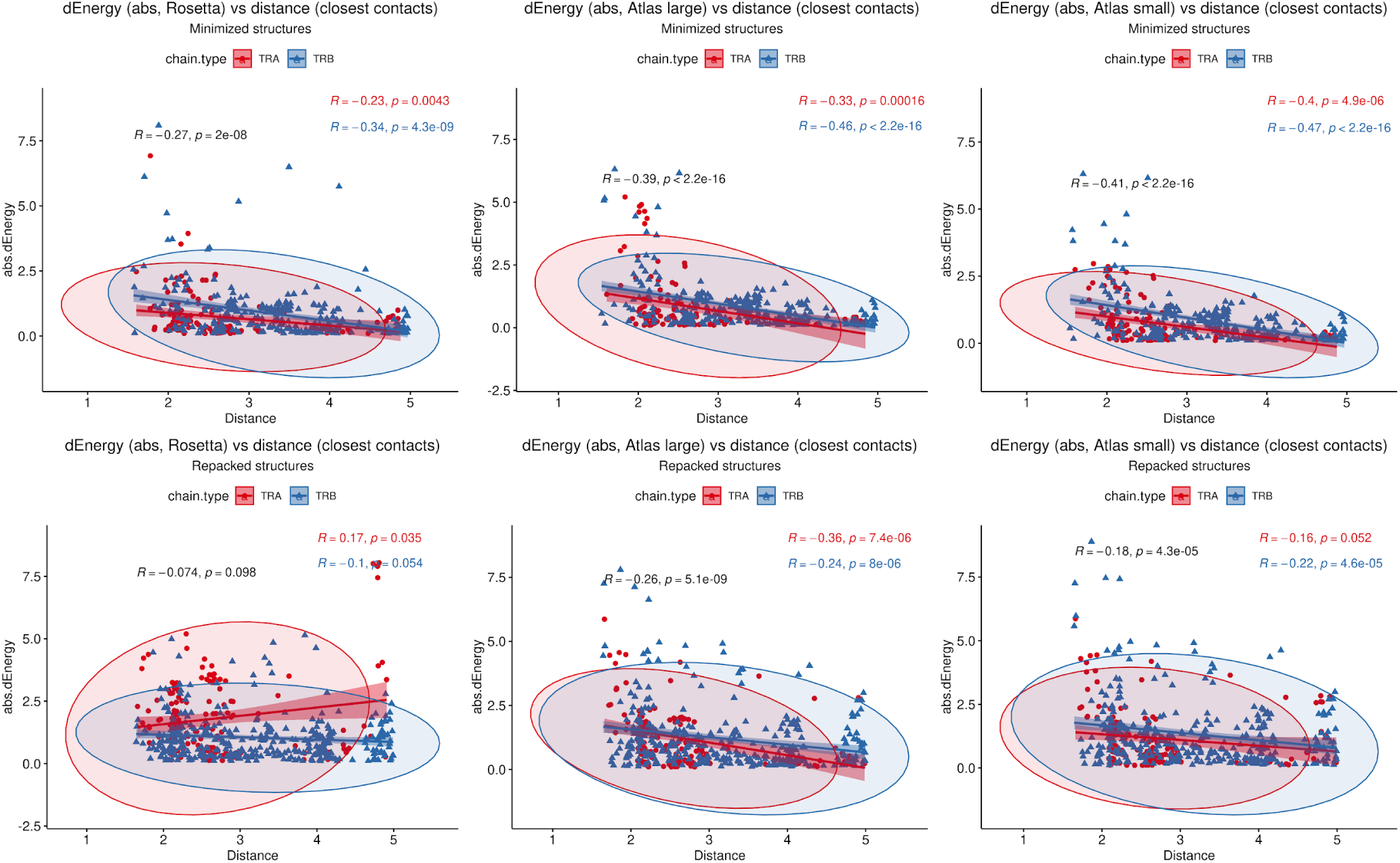
Correlation between the remoteness of contacting residues in CDR3-peptide interface and absolute dEnergy values. Overall correlation values are presented in black, and linear fittings calculated for TCR-alpha and TCR-beta chains independently are presented in red and blue respectively.

#### Correlation between physicochemical properties of cognate CDR3 and epitope residues

Using “Peptides” and “HDMD” R packages, seven groups of amino acid descriptors were calculated for the contacting residues in CDR3 loops and peptides in original and mutated TCR-pMHC complex structures. It should be mentioned that only mutated positions were taken into account. The descriptor groups were as follows: BLOSUM indices (BLOS1-BLOS10), Kidera factors (*KF1-KF13*), VHSE (*VHSE1-VHSE 8*), Cruciani properties (*PP1-PP3*), zScales (*Z1-Z5*), FASGAI (*F1-F6*) and Atchley factors (PAH, PSS, MS, CC, EC) (*32*).

The majority of calculated indices can be grouped according to the specific physicochemical features they represent: hydrophobicity, steric properties, electronic, and secondary structure features (**Table 4**).

**Table 4.**
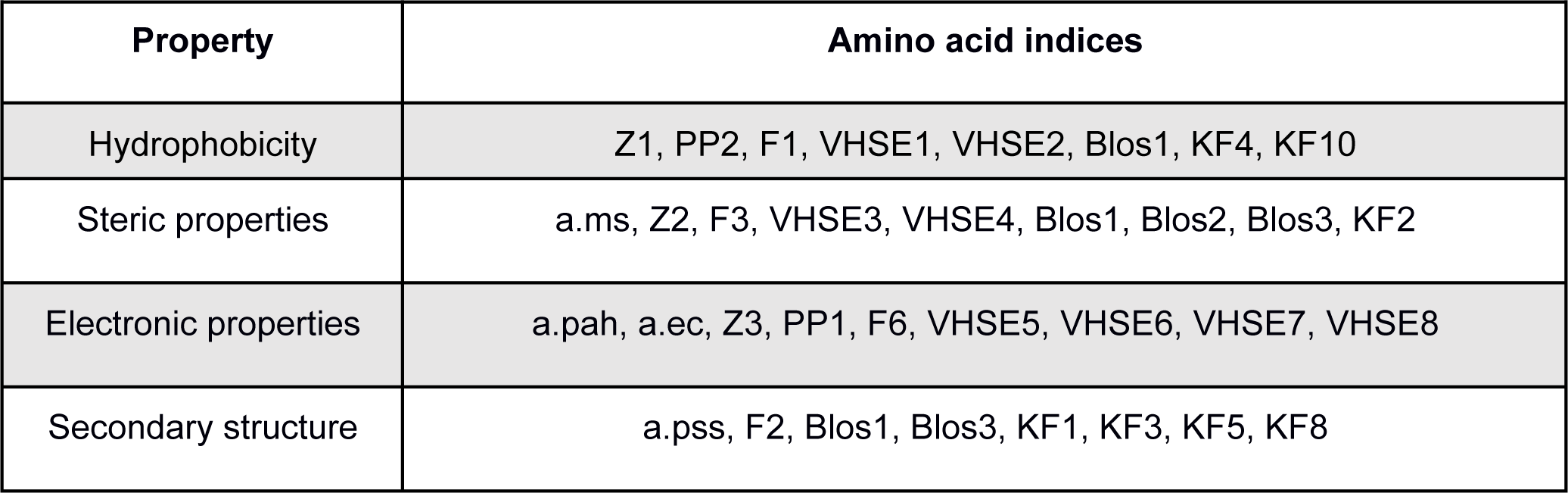
Groups of studied amino acid descriptors *Names of the Atchley indices were shortened using “a.” prefix*

The rest of the descriptors, for example, BLOS4 and BLOS9 correlate with several properties and the specific one cannot be prioritized.

To analyze the consistency between properties of contacting residues in CDR3 loops and epitopes, we performed rank correlation analysis of the corresponding factors and indices. Spearman correlations were calculated within selected groups of descriptors for all interacting residues, the closest contacting residues, and amino acid pairs interacting through at least one side chain group. The most valuable correlations in groups of descriptors were identified using the Shapiro-Wilk test. Paired correlations and their distribution for selected groups of indices are presented in **Figure 6**, and best correlations are listed in **Table 5**.

**Figure 6.**
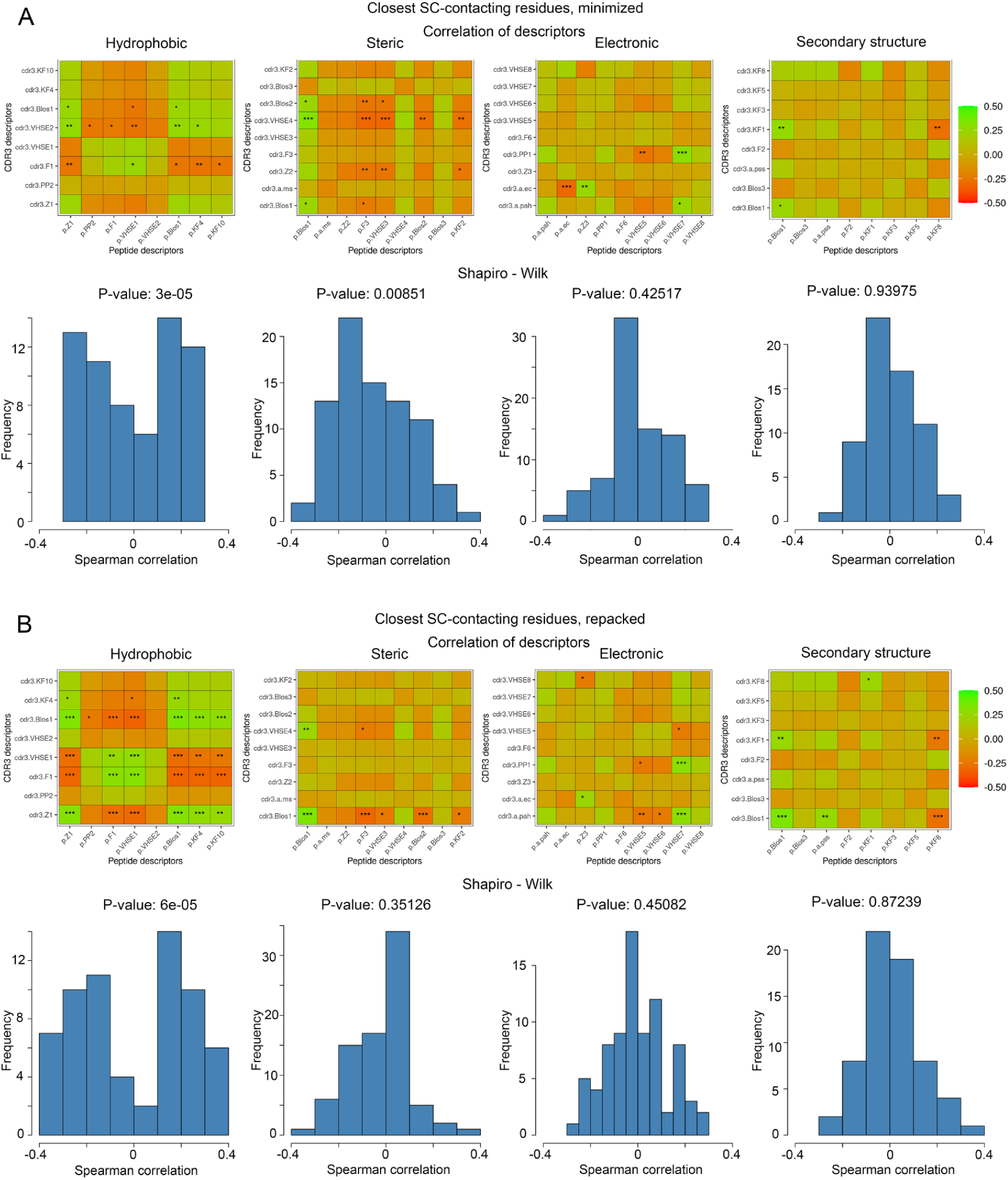
Correlations of calculated descriptors between contacting residues of CDR3 loops and epitopes. The presented correlation analysis was performed for the closest contacting residues, interacting through at least one side-chain group in A) minimized and B) repacked models. Paired correlation values are represented as waffle plots, and their distribution in groups is shown as histograms. Statistical significance of correlation is labeled with ”*”, “**” and “***” for P-values < 0.05, 0.01 and 0.001 respectively.

**Table 5.**
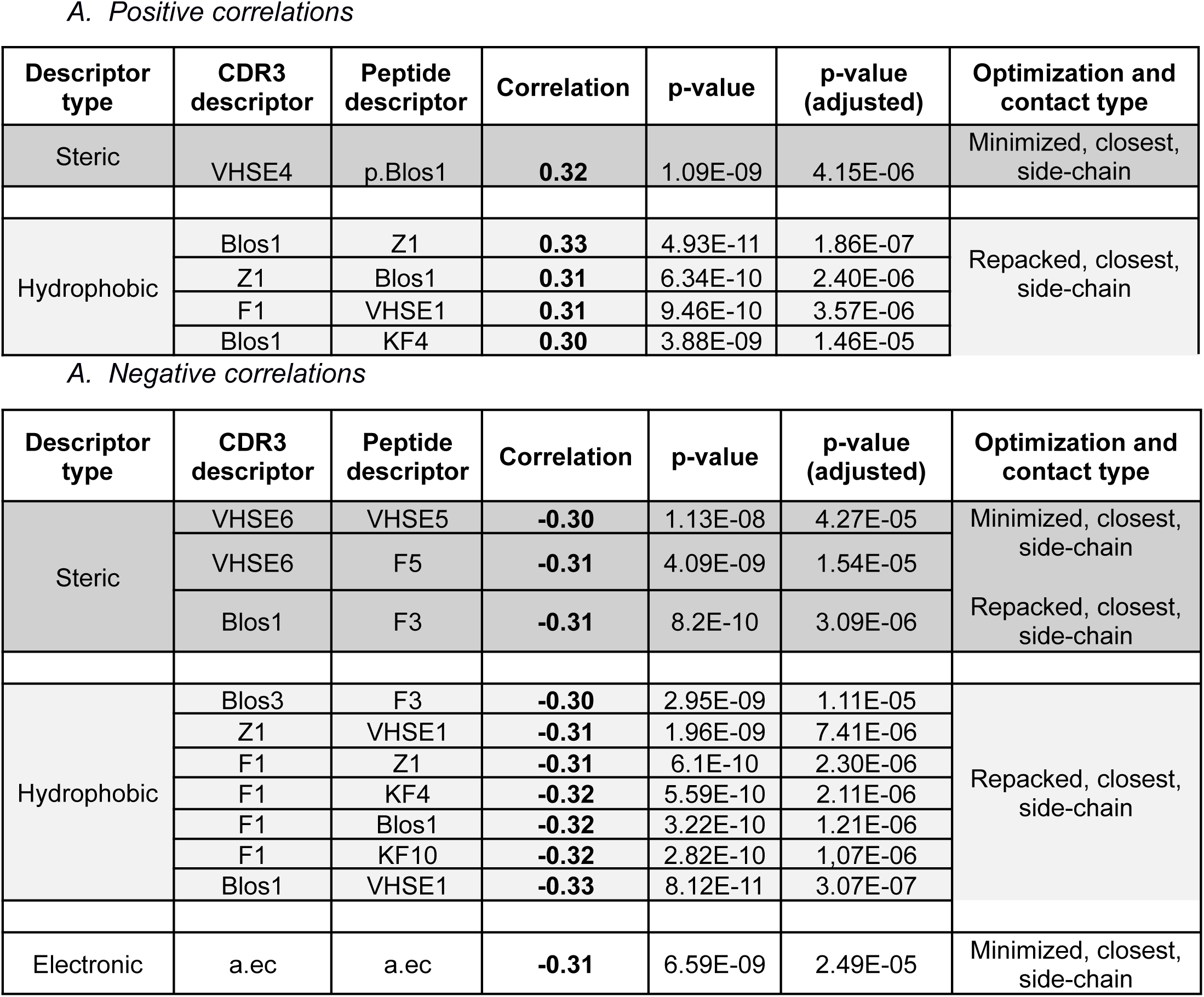
Top correlation values between calculated CDR3 and epitope residues indices.

The best correlation values were calculated for the hydrophobic and steric groups of descriptors in minimized structures, and their distribution was non-normal, according to the Shapiro–Wilk test (p-values were 3×10^-5^ and 0.0085, respectively). After repacking of modeled structures, which led to structural fitting of contacting residues and displacement of their side-chain groups, the correlations of steric descriptors became weaker, while the correlations of hydrophobic descriptors still remained highly significant (p-value = 6×10^-5^).

### Identifying residue binding features in CDR3:peptide interface using machine learning

At the final stage we performed RFE (Recursive Feature Elimination) analysis and applied the random forest (RF) algorithm to make predictive models for the affinity values of contacting amino acid residue pairs in CDR3 loops and peptides. To include a wider range of valid pairwise contacts between CDR3 loops and TCRs, we calculated amino acid descriptors and energy values for known TCR-pMHC complexes available in the PDB database. In total, we added 152 structures of human T-cell receptors with peptides and MHC-I triple complexes to the analysis.

As our modeled structures were optimized using energy minimization or repacking of contacting residues, corresponding structures were treated separately and combined with selected resolved structures from the PDB database. The two extended datasets, containing 94 amino acid descriptors and affinity values (per-residue energies) for all the studied contacting residues were divided into training and testing sets in 70% to 30% proportion. Affinity values were converted to 5 groups (factors) using quantile values.

Discretizing the target variables is a common step in machine learning procedures such as Random Forest, and there are several reasons for doing so. First, dividing the affinity values into meaningful groups can make it easier to interpret the results. Second, discretizing the values can help mitigate the effects of outliers and noise in the data. Finally, it can be useful for classification tasks, such as predicting whether a given interaction is “strong” or “weak”.

During RFE analysis of the extended datasets containing either minimized or repacked modeled structures, we identified 10 and 13 variables, respectively, as the most important predictors of the affinity values. These descriptors are presented in **Table 6**

**Table 6.** Selected amino acid descriptors during Random Feature Elimination procedure. *The top-5 descriptors according to the calculated importance are highlighted in bold font. The properties represented by the descriptors are labeled as follows: H- hydrophobic, E - electronic, S- steric, SS - secondary structure*

| Minimized + original | Repacked + original |
| --- | --- |
| **p.F1 (H)** | **p.F1 (H)** |
| **cdr3.VHSE1 (H)** | **cdr3.VHSE1 (H)** |
| **cdr3.Z1 (H)** | **cdr3.Z1 (H)** |
| cdr3.KF10 (H) | **cdr3.KF10 (H)** |
| **cdr3.KF4 (H)** | cdr3.KF4 (H) |
| cdr3.VHSE7 (E) | cdr3.VHSE7 (E) |
| cdr3.Blos8 | cdr3.VHSE8 (E) |
| **cdr3.a.ms (S)** | cdr3.a.ms (S) |
| p.a.cc | cdr3.F2 (SS) |
| cdr3.KF7 | **cdr3.KF7** |
|  | cdr3.Blos7 |
|  | cdr3.F5 |
|  | cdr3.Blos10 |

Upon analyzing the selected descriptors, it is noticeable that most of them belonged to amino acids from CDR3 loops, and the majority of the top 5 descriptors were related to the hydrophobic properties of the studied residues.

Evaluating the models built for the two selected datasets using testing data showed that the quality of their predictions was relatively similar for repacked and minimized structures. The accuracy of the model built using minimized structures was 0.4212 with a 95% confidence interval of 0.3905 - 0.4524, and the accuracy of the model using repacked structures was 0.448 with a 95% confidence interval of 0.417 - 0.4793. The Kappa statistic for the “minimized” model was 0.2765, and for the “repacked” model, it was 0.3098. The slightly higher accuracy of the model built using the extended data set of repacked structures and the higher Kappa statistic indicates better agreement between predicted and actual values compared to the second model built using minimized structures. The McNemar’s Test P-Value was also lower for the “repacked” model, suggesting that it may be a more reliable model.

## METHODS

### Identification of CDR3 sequences clusters

Clustering of CDR3 sequences from the T-cell receptor sequences database VDJdb (6) was performed according to three main rules. First, a “core” of a cluster was designated by one of the CDR3 sequences presented in a PDB structure. Second, new CDR3 sequences (neighbors) were placed in the cluster if they could be generated step-by-step introducing single amino acid substitutions one at a time. The CDR3 sequences should have the same lengths, so no deletions or insertions were allowed. Additionally, the V- and J- genes of corresponding TCRs should be the same as the one carrying the “core” CDR3 loop. Pairing of TCRα - TCRβ chains was not taken into account. Third, the Hamming distance between the core and each of the neighbor CDR3 sequences in the cluster was limited to 3. Thus, the mutation path from the CDR3 sequence from the PDB database and the most diverse sequence in the cluster should consist of not more than 3 consequent single mutations. The validation of our proposed modeling approach and the explanation of why we selected a limit of 3 mutations is discussed in **Supplementary Note 1**.

### Step-by-step modeling of CDR3 loops

A Rosetta package (35) was used to implement an *in silico* single amino acid mutation modeling approach. The Rosettascripts mover “MutateResidue” was used to replace selected residues while retaining main chain heavy atom positions according to the initial template structures, modifying only side chain atoms.

Further structural and energy optimization was performed using the relax protocol with additional restraints on atom coordinates, L-BFGS minimization algorithm and *ref2015* score function. Additionally, the repacking protocol of Rosettascripts was used to optimize contacting residues in modeled TCR – peptide - MHC complexes. Repacking was performed for CDR3 loop and peptide residues situated closer than 5Å from each other, as these residues were estimated as contacting in modeled complexes.

### Structural data analysis

Identification of close CDR3 and peptide residues in modeled complexes was performed using in house Java and Bash scripts. Contacting parts of residues (side and main chains) were identified using Residue Interaction Network Generator approach (RING) (36).

Calculation of energy values of contacting TCR-pMHC interfaces in modeled structures, as well as *RMSD* values between modeled and original CDR3 loop structures was performed using Rosetta score_jd2 application. Energy values were calculated using three variations of Rosetta ref2015 scoring function: “full” ref2015 energy function with all default energy terms included, and two “limited” variations, set by lists of selected terms. The first preset (“Large patch”) included *fa_atr* (attractive portion of the Lennard Jones potential), *fa_sol* (Lazaridis-Karplus solvation energy), *hbond_sr_bb* (h-bond energy, short-range backbone-backbone), *hbond_lr_bb* (bond energy, long-range backbone-backbone), *hbond_bb_sc* (h-bond energy, backbone-sidechain) and *hbond_sc* (h-bond energy, sidechain-sidechain) energy terms. The second (“Small patch”) consisted of the same terms as the “Large patch”, except for *hbond_sr_bb* and *hbond_bb_sc*.

Amino acid descriptors and indices were calculated using “Peptides” (BLOSUM similarity matrix indices, Kidera factors, VHSE, Cruciani properties, zScales, FASGAI) and “HDMD” (Atchley factors) packages in R (37,38).

### Statistical analysis and machine learning

Statistical analysis of computed energy values of the modeled structures and properties of contacting amino acid residues in TCR-pMHC complexes was performed using R packages. RFE analysis was performed on the training set consisting of 94 amino acid descriptors, calculated for contacting amino acid residues in CDR3 loops of TCRs and peptides to select the most important descriptors for predicting the target variable. Random forest algorithm was then used to train a model, which was used to make predictions on the testing set. Per-residue energy values were grouped into 5 clusters and used as target variables during the RFE analysis and RF modeling. The performance of the model was evaluated using confusion matrix analysis. Calculations and data preparation were performed using Caret and RandomForest packages in R.

### Data and code availability

Modeled and template structures are available at Zenodo (https://doi.org/10.5281/zenodo.7845844). Custom R scripts used in this study are available at GitHub (https://github.com/antigenomics/vdjdb-structure).

## Discussion

In this study we proposed an *in silico* approach for template-based modeling of highly similar CDR3 loops in full TCR-peptide–MHC complexes using stepwise introduction of single amino acid substitutions. Using our approach, we modeled 1585 structures based on 29 templates and studied the non-redundant set of structures by grouping identical TCR alpha and beta chains and utilized the dataset to study the determinants of antigen recognition on a single-residue level. Our results greatly extend the number of available structures with different hypervariable CDR3 sequence variants, including 16 synthetic TCR alpha structures and 38 TCR beta structures modeled based on 7RTR and 7N1F SARS-CoV-2 antigen-TCR complex templates.

Upon analyzing these structures, we observed that widely used standard BLOSUM62 and BLOSUM100 scores have little correlation with the impact of amino acid mutations in CDR3 loops on TCR–epitope binding energy as long as those substitutions appear in non-contacting parts of loops. This finding suggests that the analysis and comparison of CDR3 sequences in context of epitope recognition should mainly consider contacting residues, and conventional full-length sequence alignment alone is not sufficient to compare binding affinity of CDR3 sequences to the same antigen.

Also, we found that contacting residues in CDR3b can have a greater effect on TCR–peptide recognition than those in CDR3a loops. This conclusion is based on the analysis of the impact of CDR3 amino acid variations on interaction energy values, depending on their remoteness from peptides and correlation between BLOSUM indices and calculated absolute values of dEnergy of interacting interfaces. In both cases corresponding properties and values of CDR3b residues showed better correlations in comparison to CDR3a.

Analysis of physicochemical properties of contacting residues in CDR3 loops and epitopes, described by different descriptors, showed that hydrophobicity may be an important factor governing recognition and affinity of binding between CDR3 loops and epitopes in TCR-pMHC complexes, yet in all analysis correlation values were moderate. This is in agreement with the fact that all studied amino acid residue substitutions have limited impact on binding of CDR3 loops to peptides in order to preserve T-cell receptor’s recognition.

The pipeline described in this paper can be reused when more templates and/or TCR specificity data becomes available, greatly extending the amount of available structural data on antigen recognition by TCRs. Possible extensions to this work include models in which the antigen is substituted by a similar peptide that binds the same MHC in order to monitor for potential TCR cross-reactivity and perform in-depth study of the determinants of TCR specificity using molecular dynamics.

## Acknowledgements

This work was supported by grant № 075-15-2019-1789 from the Ministry of Science and Higher Education of the Russian Federation.

## Supplementary Information

**Figure S1.**
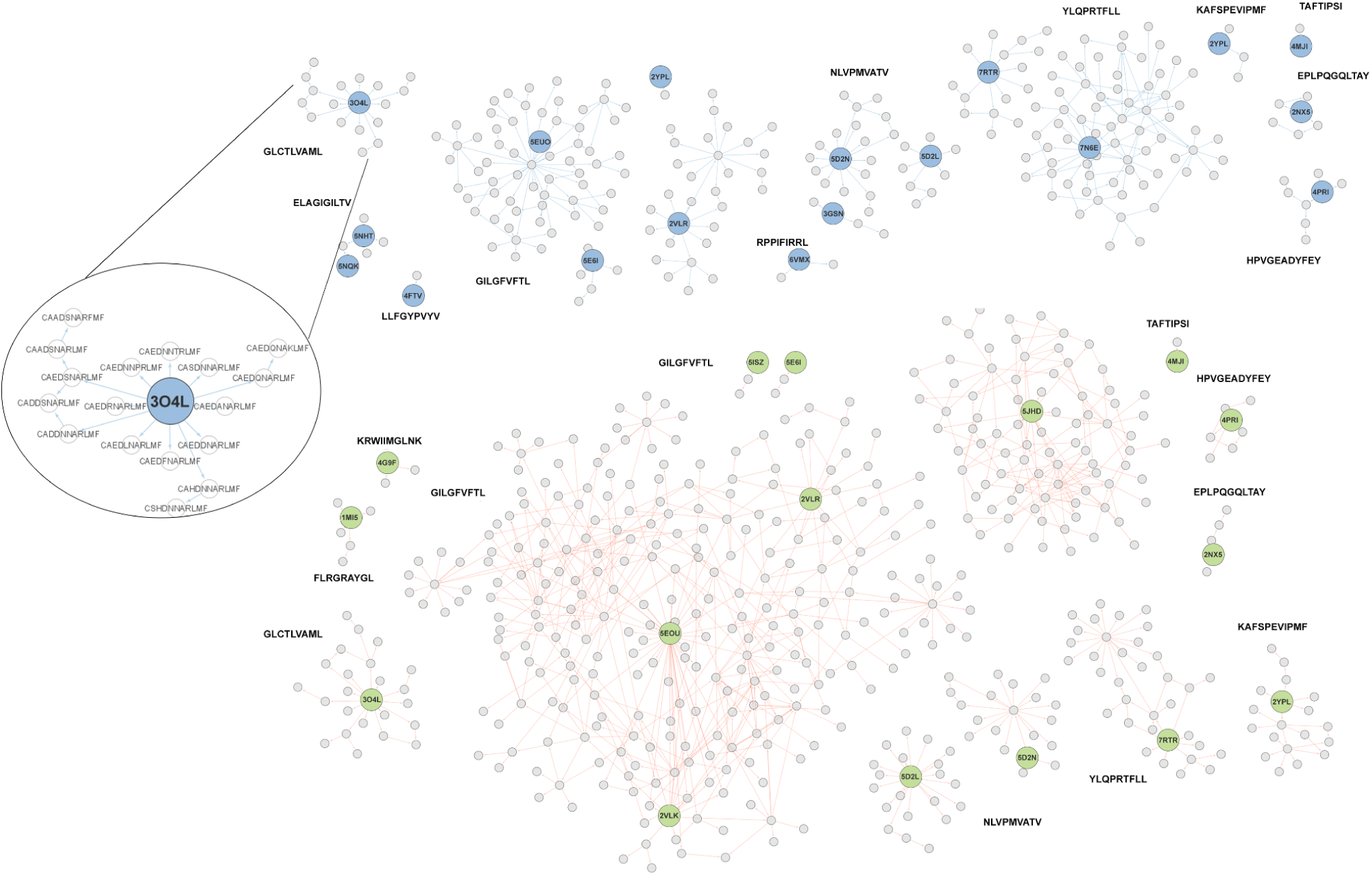
Graph with paths used for stepwise modeling of CDR3 loops. CDR3α sequences are connected with blue edges and sequences of CDR3β – red. Initial templates are presented as circles with PDB identifiers and are colored blue in CDR3α and green in CDR3β clusters. Epitope labels are placed near corresponding connected components.

**Figure S2.**
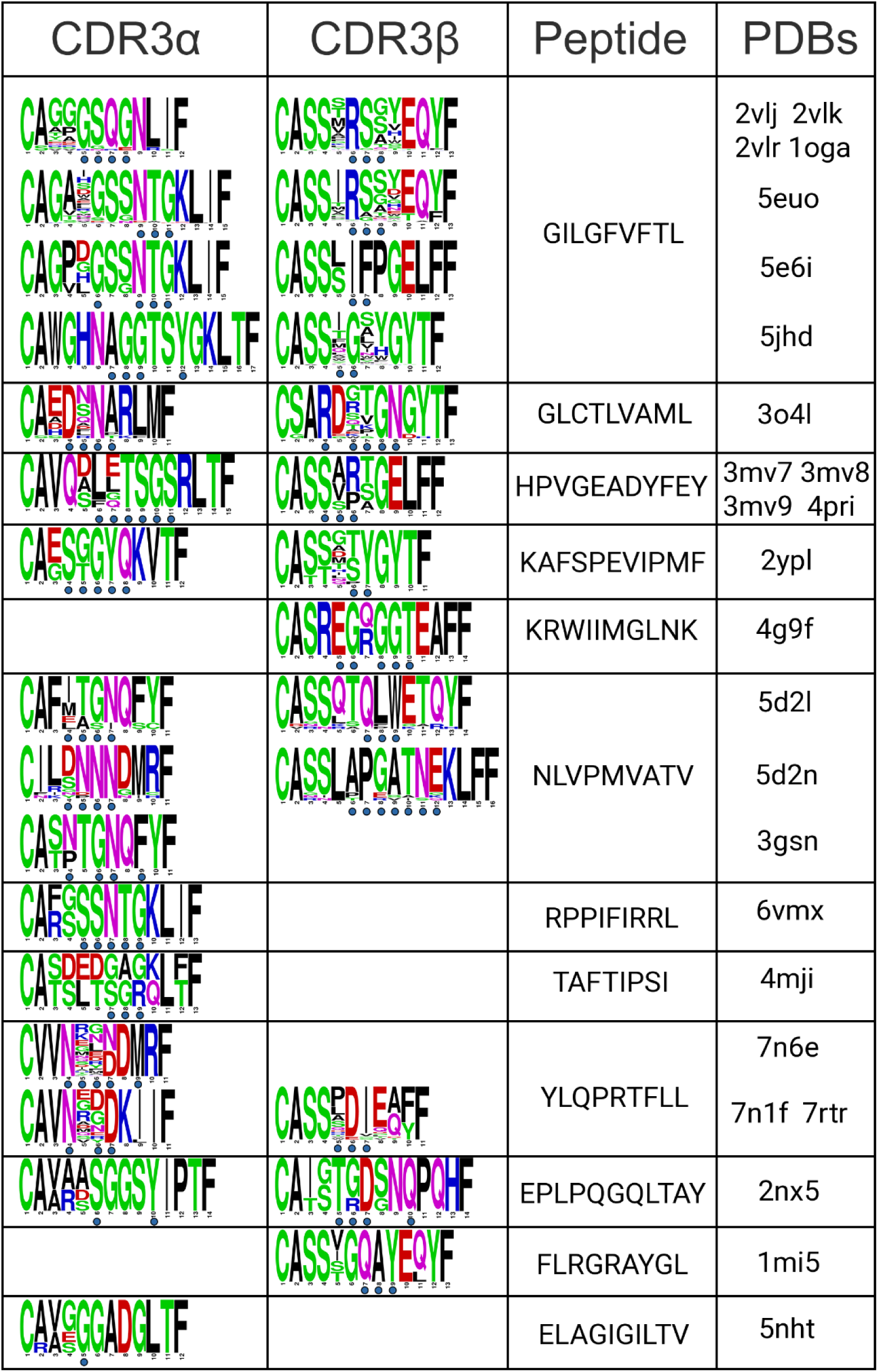
Logo-formatted sequences of CDR3 loops in modeled structures. Residues, contacting with peptides are marked with blue circles.

**Table S1.**
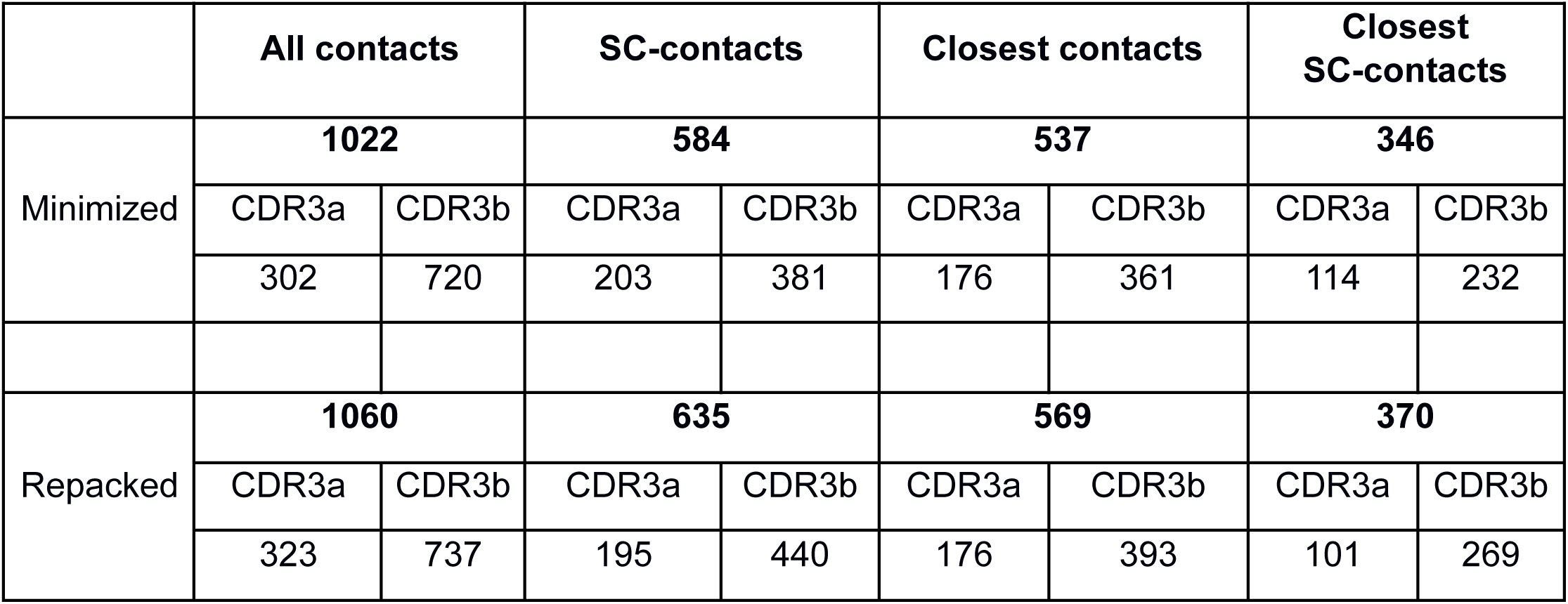
Number of paired CDR3 – epitope contacting residues in modeled structures. “All contacts”- all contacting pairs formed by mutated residue of CDR3 loops with epitope amino acids, situated closer than 5Å apart. “SC- contacting”- contacting pairs formed by mutated residue of CDR3 loops with epitope amino acids, situated closer than 5Å apart. At least one residue in the pair interacts by its side chain group. “Closest contacts” – only the closest residue of epitope selected for each mutated residue of CDR3 loop in each modeled structure. “Closest SC-contacts” - only the closest, interacting by at least one side chain group, residues of epitope selected for each mutated residue of CDR3 loop in each modeled structure.

## Supplementary Note 1

### Validation of the proposed stepwise single amino acid substituting modeling approach

#### Materials and methods

##### TCR-pMHC homology modeling

Modeling of the selected TCR structures was performed using our approach, TCR-pMHC modeling (*11*) service and TCRModel algorithm (*10*) implemented in the Rosetta package. The models were selected based on their sequence similarity to known templates from the PDB database. CDR3 loops were modeled by introducing less than 6 amino acid substitutions starting from the known template, and the resulting “target” TCR chain should also be resolved in the PDB database. During modeling using TCRModel, target structures were eliminated from the templates library. TCR-pMHC modeling was performed using the online service with default settings (http://www.cbs.dtu.dk/services/TCRpMHCmodels/index.php). These models were used to validate our modeling and optimization approaches.

In addition to energy minimization and repacking of contacting residues, loop refinement was performed for our models using a kinematic protocol implemented in the LoopRemodel Rosettascripts mover. During the refinement procedure the four terminal residues (two residues at the N-terminus and two - at C-terminus) of the CDR3 loop were fixed, while the rest of the backbone and side chains were allowed to be flexible.

##### Molecular dynamics

Molecular dynamics simulations were carried out using Gromacs-2020.5 software package (*37*). The systems were neutralized by adding Na^+^ and Cl^-^, and solvated with TIP3P water models. MD calculations were performed using the Amber99-SB force field. Energy minimization was sequentially performed in vacuum and in solvation for 150,000 steps. Subsequently, temperature was increased to 300K and pressure was set to 1 atm using NVT and NPT ensembles, respectively. Molecular dynamics simulations were performed on 50 ns trajectories with 2 fs time steps.

#### Results

##### Minimization vs. repacking of TCR-pMHC contacting area

Initial visual inspection of some previously modeled complexes using our approach revealed sterical issues that can occur during spatial positioning of large amino acid residues instead of small ones, and their orientation was altered depending on the final optimization step. For example, finalizing repacking optimization led to the misplacement of Arg and Tyr side chain groups in mutated β-chain CDR3 loop from 2VLR PDB structure (**Figure SN1**) compared to a structure received after energy minimization only. To validate the stability of these modeled structures, we performed molecular dynamics calculations for three selected pMHC-TCR complexes: two modeled structures with the “CASSIRWAYEQYF” CDR3 loop of the beta-chain of TCR, built based on 2VLR PDB structures with just energy minimization or repacking of contacting residues as the final optimization step; and the original 2VLR PDB structure that was used as an initial template for the modeling process.

During the analysis of the MD trajectories, the most attention was paid to the displacement of peptide-MHC complex with respect to TCR since the observed displacement of side chain groups in contacting residues might influence the stability of the TCR-pMHC interaction. It was shown that in the original TCR-pMHC complex structure (PDB id: 2VLR) the pMHC position was the most stable (**Fig.SN1D**). RMSD values for the peptide-MHC molecules were in the range of 3.5-5A during molecular dynamics.

On the other hand, the modeled complex that was built using only energy minimization as the final optimization step was unstable (**Figure SN1D**), and the MHC molecule with the peptide dramatically shifted from their initial position during MD (**Figure SN1B**). Such low affinity of this modeled complex might be explained by a bad structural “complementary” in the TCR-pMHC contacting interface due to the unfavorable positioning of replaced side chain groups.

Changes in RMSD values for the MHC-peptide molecules in the third (repacked) complex showed that at the beginning of molecular dynamics, the pMHC complex shifted from its initial pose and returned back after 15 - 20 ns of MD.The alteration of MHC-TCR docking pose is shown in **Figure SN1C**.

These results show that repacking of contacting residues might be a necessary step while modeling CDR3 loops in TCR-pMHC complexes using our step-by-step single mutation approach to achieve more stable complexes and cover larger conformational space of TCR-pMHC contacting interfaces.

**Figure SN1.**
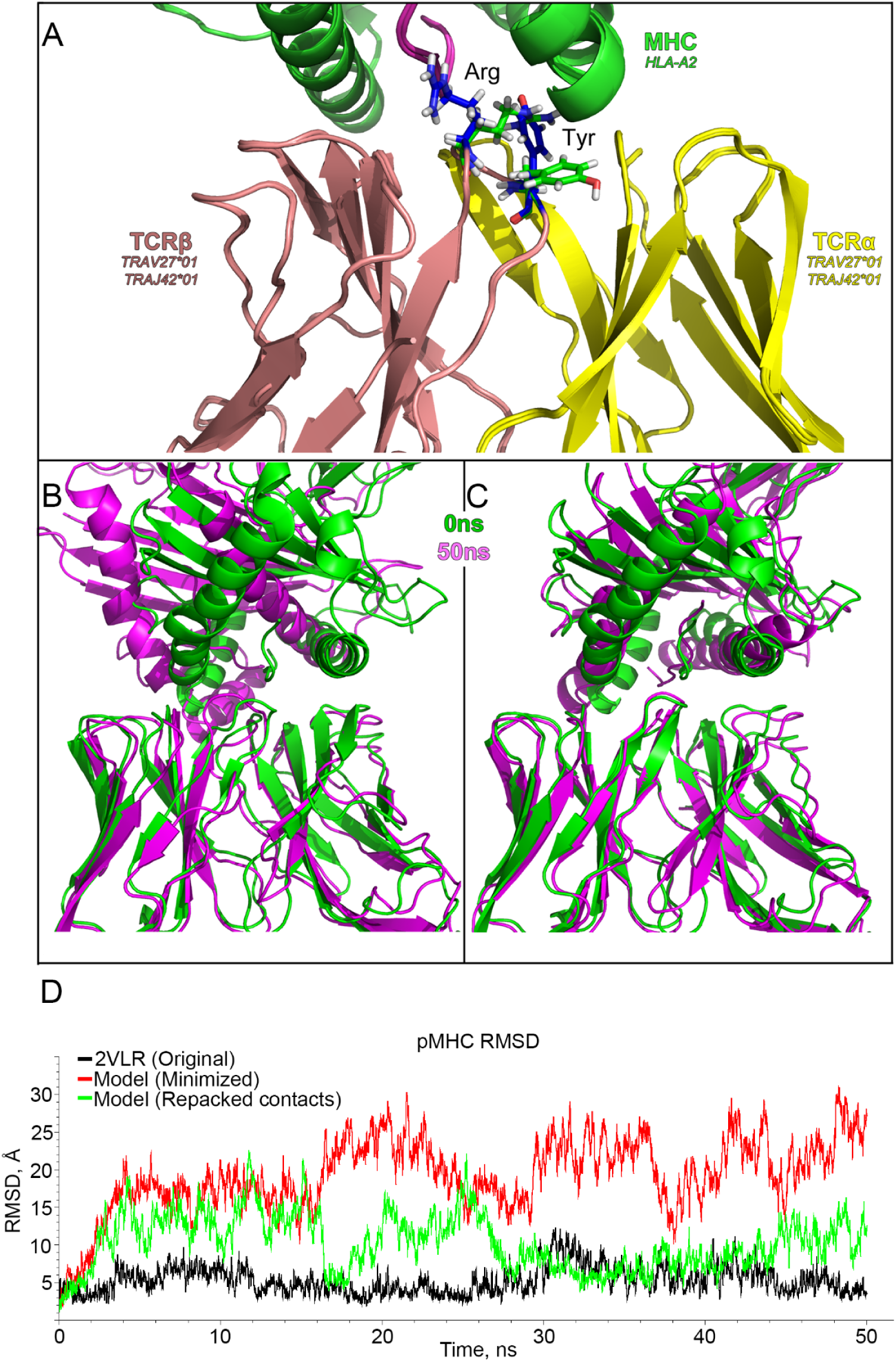
Stability of the original and modeled TCR-pMHC complexes and structural comparison of CDR3 loops after energy minimization or repacking of contacting residues. A) Spatial displacement of Arg and Tyr side chain groups after repacking of contacting residues in pMHC – TCR complex built from 2VLR. Repacked amino acids are colored blue, mutated and just minimized structures are green. B) Displacement of the peptide-MHC complex during molecular dynamics of just minimized modeled structures and C) after repacking of the contacting interface. D) RMSD values, calculated for peptide-MHC complexes during molecular dynamics. Values of the original PDB structure (2VLR) are represented as a green line, minimized modeled structure - as black, and repacked modeled structure - as red. All structures of complexes were aligned by TCR chains.

##### Validation of modeling approaches using reference PDB structures

To validate the proposed mutation modeling approach and compare our results with corresponding models built using available TCR modeling services, we analyzed sequences of all CDR3 loops of ternary TCR-pMHC complexes available in the PDB database. All found sequences were pairwise aligned separately for TCR-alpha and TCR-beta chains. Analyzing the built alignments, we selected 27 CDR3 sequences that could be modeled by introducing up to 6 sequential substitutions starting from some other template structures presented in the PDB database. Hereinafter, the number of introduced amino acid substitutions will be referred to as the “mutation order”. The list of all selected CDR3 sequences with initial templates for the modeling process and their original PDB structures is presented in **Table SN1**

**Table SN1.** Structures, used for validation of modeling approaches.

| PDB template | TCR | CDR3 | PDB original | Order |
| --- | --- | --- | --- | --- |
| 3qeq | TRA | CAFITGNQFYF | 5d2l | 2 |
| 5e6i | TRA | CAGAIGPSNTGKLIF | 5euo | 3 |
| 3qdm | TRA | CARNTGNQFYF | 3gsn | 2 |
| 1ao7 | TRB | CASRPGLMSAQPEQYF | 4ftv | 4 |
| 5yxn | TRB | CASRRGSAELYF | 5yxu | 3 |
| 2vlr | TRB | CASSDTVSYEQYF | 5nme, 5nmf, 5nmg | 3 |
| 5nmg | TRB | CASSDWVSYEQYF | 1lp9, 2j8u, 2jcc,<br>2uwe | 1 |
| 6bj2 | TRB | CASSFNMATGQYF | 5brz, 5bs0 | 5 |
| 1oga | TRB | CASSIRSSYEQYF | 5euo, 5hho | 1 |
| 2vlj | TRB | CASSIRSSYEQYF | 5euo, 5hho | 1 |
| 5euo | TRB | CASSIRSSYEQYF | 5hho | 1 |
| 4l3e | TRB | CASSLSFGTEAFF | 3qdg, 3qdj, 6am5,<br>6amu | 1 |
| 5wkf | TRB | CASSPVTGGIYGTYF | 3gsn | 5 |
| 6avf | TRB | CASSQTLWETQYF | 5d2l | 4 |
| 2vlj | TRB | CASSSRASYEQYF | 2vlr | 1 |
| 2vlr | TRB | CASSSRSSYEQYF | 1oga, 2vlj, 2vlk,<br>2xn9, 4c56, 5hhm | 1 |
| 5euo | TRB | CASSSRSSYEQYF | 1oga, 2vlj, 2vlk,<br>2xn9, 4c56, 5hhm | 1 |
| 4mji | TRB | CASSSWDTGELFF | 3vxr, 3vxs | 4 |
| 3qdg | TRB | CASSWSFGTEAFF | 4l3e, 6d78, 6dkp | 1 |
| 3vxr | TRB | CASSWSFGTEAFF | 4l3e, 6d78, 6dkp | 6 |
| 6amu | TRB | CASSWSFGTEAFF | 4l3e, 6d78, 6dkp | 1 |
| 2f53 | TRB | CASSYLGNTGELFF | 2p5e, 2p5w, 2pye | 1 |
| 2p5w | TRB | CASSYVGNTGELFF | 2bnq, 2bnr, 2f53,<br>2f54, 6q3s | 1 |
| 4mnq | TRA | CAVDSATSGTYKYIF | 4c56, 5men | 4 |
| 5w1v | TRA | CAVLSSGGSNYKLTF | 5eu6 | 4 |
| 5nqk | TRA | CAVSGGYQKVTF | 2ypl | 6 |
| 4g9f | TRA | CLVGEILDNFNKFYF | 6avf | 6 |

All T-cell receptor structures containing the selected CDR3 sequences were modeled using the TCRModel package of Rosetta software (10), TCR-pMHC modeling service (11) and our mutation approach. In addition to energy minimization and repacking of contacting residues, we performed a loop refinement procedure in our modeling approach as the final optimization step. Refinement was performed using Rosetta “LoopRemodel” mover with four “fixed” residues: two at each end of CDR3 loops.

As there can be some variability during conformational search and optimization of models, several runs of modeling were performed for each of the studied approaches: two runs of modeling were performed for our approach with different optimization procedures, two for the TCR-pMHC modeling service and three runs for TCRmodel.

RMSD values between compared structures were calculated using the Profit application (Martin, A.C.R. and Porter, C.T., http://www.bioinf.org.uk/software/profit/, (38)) separately for all atoms of CDR3 loops and for C-alpha atoms only. During the study all models were superposed with original (target) PDBs by corresponding TCR chains or by CDR3 loops. Thus, RMSD values, calculated for structures aligned by CDR3 loops, represent the “accuracy” of predictions of corresponding loop shapes, while alignment using TCR chains allows the evaluation of both the loop shapes and their orientation. Results are presented in **Figure SN2**.

If modeled structures constructed based on one template had several known original target structures, mean values were calculated for the corresponding deviations. RMSD values of different modeling runs are presented independently on the graph

**Figure SN2.**
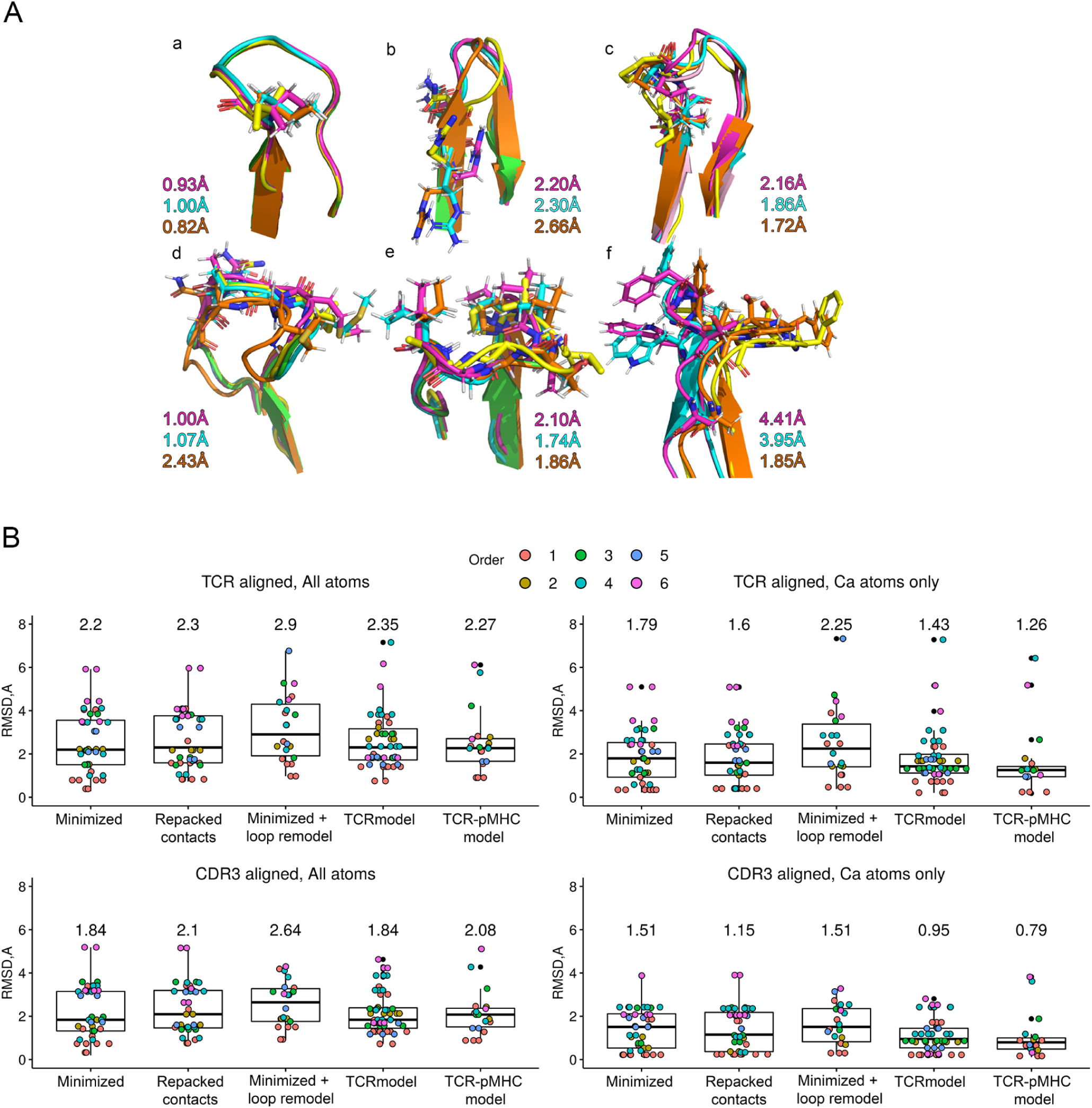
Comparison of modeled CDR3 structures with corresponding known conformations of CDR3 loops from PDB data bank. A) Structural comparison of models with different mutation orders. RMSD values, calculated for all atoms of CDR3 loops in structures aligned by TCR chains, are presented for three modeling approaches: our approach with energy minimization (magenta), our approach with repacking of contacting residues (cyan) and TCRmodel (orange). Initial template structures are colored green and original structures with target CDR3 loops are colored yellow. CDR3 sequences and mutation orders are as follows: a) Mutation order = 1, CDR3 sequence: CASSIRSSYEQYF (TRB, original structure- 5hho, 5euo; template- 1oga), b) Mutation order = 2, CDR3 sequence: CARNTGNQFYF (TRA, original structure- 3gsn; template - 3qdm), c) Mutation order = 3, CDR3 sequence: CAGAIGPSNTGKLIF (TRA, original structure- 5euo; template - 5e6i), d) Mutation order = 4, CDR3 sequence: CASRPGLMSAQPEQYF (TRB, original structure- 4ftv; template - 1ao7), e) Mutation order = 5, CDR3 sequence: CASSPVTGGIYGYTF (TRB, original structure- 3gsn; template – 5wkf), f) Mutation order = 6, CDR3 sequence: CASSWSFGTEAFF (TRB, original structure- 4l3e; template - 3vxr) B) Boxplots of RMSD values between modeled CDR3 conformations and loops from original PDB structures. Median values for each modeling approach are presented in corresponding boxes. Models built using the mutation approach with finalizing energy minimization are labeled as “Minimized + loop remodel” and “Minimized” depending on whether loop remodel algorithm was applied or not. Models refined by repacking of all contacting residues between pMHC and TCR are labeled “Repacked contacts”, and models built using TCRModel or TCR-pMHC modeling approaches are labeled “TCRModel” and “TCR-pMHC model” respectively

Analysis of Cα atoms only in aligned CDR3 loops showed that all studied approaches allowed us to model conformations of the main chains of CDR3 loops close to the original structures. The median values for our approaches were 1.15Å for repacked structures and 1.51 Å for structures after just energy minimisation. The corresponding values for TCRmodel and TCR-pMHC modeling services were 0.95Å and 0.79Å.

Several outliers can be noted on the boxplots. These high RMSD values are associated with models carrying 6 substitutions (for all approaches) or 6-5 substitutions (for the TCR-pMHC model service).

As expected, median values of RMSD were higher during the analysis of full-atom structures of CDR3 loops including side-chain groups. The most significant increase was observed for structures modeled using the TCR-pMHC modeling service (from 0.79Å up to 2.08Å). The smallest median values were calculated for mutated structures with finalizing minimization and structures received using TCR-model (1.84Å for both mentioned approaches). CDR3 loops constructed using our approach with the loop refinement procedure at the final step had the lowest accuracy according to median RMSD values - 2.64Å during the analysis of structures aligned by CDR3 loops and 2.9Å for models aligned by corresponding TCR chains.

Analyzing all RMSD values, it can be observed that most of the modeled structures carrying from 4 up to 6 substitutions had lower accuracy than structures with mutation orders of 1-3. Median values of RMSD, calculated for structures carrying a different number of substitutions are presented in **Figure SN3**

**Figure SN3.**
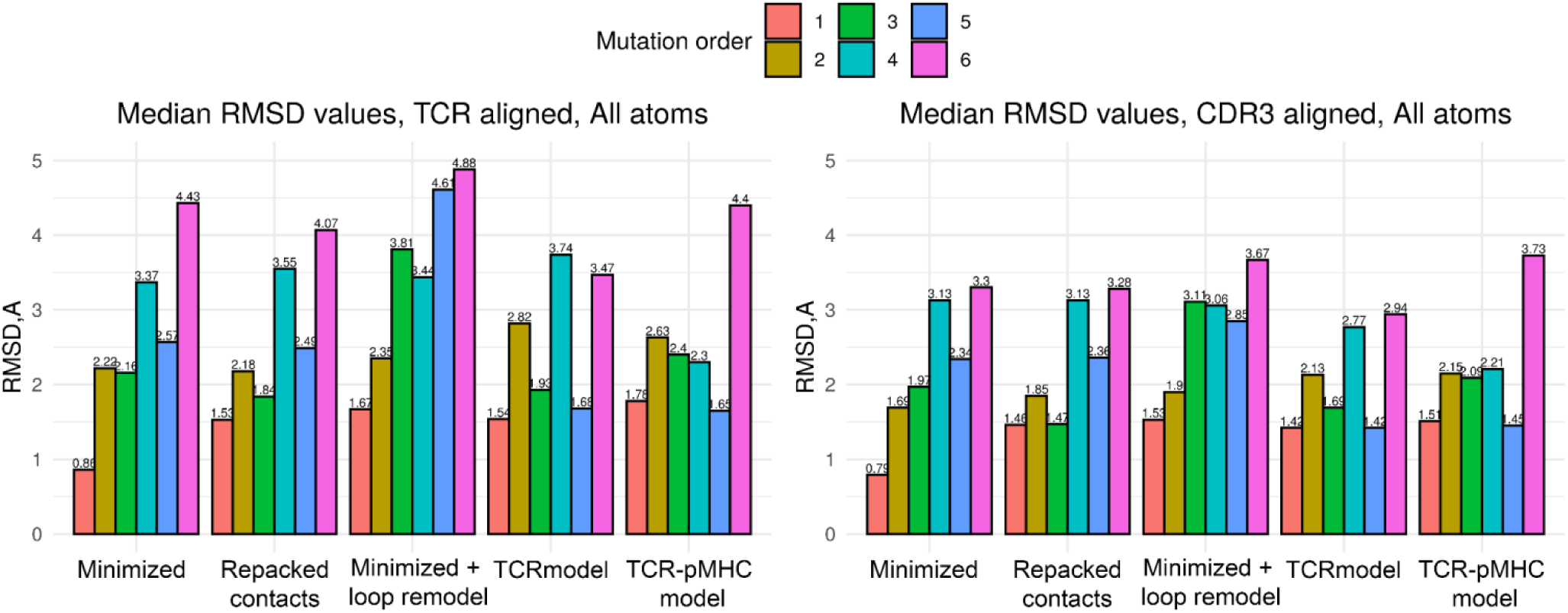
Median RMSD values, calculated for each mutation order individually.

It can be observed that the increase in RMSD values in all modeled structures correlated with the “mutation order”. According to the calculated RMSD values, the best models (closest to known structures) carried only one substitution and were built using our mutational protocol with only minimization as the final refinement step. The median values were 0.79Å and 0.86Å during the analysis of models aligned with the original structures by CDR3 loops and TCR chains, respectively. RMSD values for all other modeling approaches were in the range of 1.42Å −1.67 Å. The conformation and spatial orientation of CDR3 loops (**Fig. SN3**, TCR aligned, all atoms), carrying two substitutions, were better predicted by our approaches with median RMSD values between 2.18Å and 2.35Å. TCRmodel and TCR-pMHC modeling service predicted corresponding structures with 2.82Å and 2.63Å deviation, respectively. Third-order mutations were modeled with similar RMSD values among all tested approaches, and higher median deviation values were calculated for our approach with loop remodeling optimization.

According to median RMSD values starting from mutation order 4 and higher, the most accurate results were received using the TCR-pMHC modeling service. It could be explained by its modeling algorithm: the service uses its own templates database and there are no options to exclude target structures used in our benchmark. And, as far as we selected CDR3 loops that were available in the PDB database, TCR-pMHC models service could actually use them to perform homology modeling.

This validation step showed that our approach with finalizing energy minimization or repacking of contacting residues in TCR - pMHC interface can be used to model CDR3 loops carrying up to 3 mutations with an acceptable accuracy. However, loop refinement, on the other hand, can decrease the quality of models with 1-3 mutation order.

## References

1. La Gruta NL, Gras S, Daley SR, Thomas PG, Rossjohn J. Understanding the drivers of MHC restriction of T cell receptors. Nat Rev Immunol. 2018 Jul;18(7):467–78.

2. Schatz DG, Swanson PC. V(D)J recombination: mechanisms of initiation. Annu Rev Genet. 2011;45:167–202.

3. Goldrath AW, Bevan MJ. Selecting and maintaining a diverse T-cell repertoire. Nature. 1999;402(6759):255–62.

4. Lee CH, Salio M, Napolitani G, Ogg G, Simmons A, Koohy H. Predicting Cross-Reactivity and Antigen Specificity of T Cell Receptors. Front Immunol. 2020;11:565096.

5. Demonstrated Protocol, CG000203 [Internet]. 10x Genomics Support; 2022. Available from: https://www.10xgenomics.com/support/single-cell-immune-profiling/documentation/steps/sample-prep/cell-labeling-with-d-code-dextramer-r-for-single-cell-rna-sequencing-protocols

6. Goncharov M, Bagaev D, Shcherbinin D, Zvyagin I, Bolotin D, Thomas PG, et al. VDJdb in the pandemic era: a compendium of T cell receptors specific for SARS-CoV-2. Nat Methods. 2022 Sep;19(9):1017–9.

7. Vita R, Mahajan S, Overton JA, Dhanda SK, Martini S, Cantrell JR, et al. The Immune Epitope Database (IEDB): 2018 update. Nucleic Acids Res. 2019 Aug;47(D1):D339–43.

8. Tickotsky N, Sagiv T, Prilusky J, Shifrut E, Friedman N. McPAS-TCR: a manually curated catalogue of pathology-associated T cell receptor sequences. Bioinformatics. 2017;33(18):2924–9.

9. Gowthaman R, Pierce BG. TCR3d: The T cell receptor structural repertoire database. Bioinformatics. 2019;35(24):5323–5.

10. Berman HM, Westbrook J, Feng Z, Gilliland G, Bhat TN, Weissig H, et al. The Protein Data Bank. Nucleic Acids Res. 2000 Jan;28(1):235–42.

11. Gowthaman R, Pierce BG. TCRmodel: high resolution modeling of T cell receptors from sequence. Nucleic Acids Res. 2018 Feb;46(W1):W396–401.

12. Jensen KK, Rantos V, Jappe EC, Olsen TH, Jespersen MC, Jurtz V, et al. TCRpMHCmodels: Structural modelling of TCR-pMHC class I complexes. Sci Rep. 2019 Oct;9(1):14530.

13. Bradley P. Structure-based prediction of T cell receptor:peptide-MHC interactions. eLife [Internet]. 2023;12. Available from: http://dx.doi.org/10.7554/eLife.82813

14. Schritt D, Li S, Rozewicki J, Katoh K, Yamashita K, Volkmuth W, et al. Repertoire Builder: high-throughput structural modeling of B and T cell receptors. Mol Syst Des Eng. 2019 May;4(4):761–8.

15. Klausen MS, Anderson MV, Jespersen MC, Nielsen M, Marcatili P. LYRA, a webserver for lymphocyte receptor structural modeling. Nucleic Acids Res. 2015 Jan;43(W1):W349–55.

16. Reynisson B, Alvarez B, Paul S, Peters B, Nielsen M. NetMHCpan-4.1 and NetMHCIIpan-4.0: improved predictions of MHC antigen presentation by concurrent motif deconvolution and integration of MS MHC eluted ligand data. Nucleic Acids Res. 2020 Feb;48(W1):W449–54.

17. Li S, Wilamowski J, Teraguchi S, van Eerden FJ, Rozewicki J, Davila A, et al. Structural Modeling of Lymphocyte Receptors and Their Antigens. Methods Mol Biol. 2019;2048:207–29.

18. Grazioli F, Mösch A, Machart P, Li K, Alqassem I, O’Donnell TJ, et al. On TCR binding predictors failing to generalize to unseen peptides. Front Immunol. 2022;13:1014256.

19. Dhusia K, Su Z, Wu Y. A structural-based machine learning method to classify binding affinities between TCR and peptide-MHC complexes. Mol Immunol. 2021 Nov;139:76–86.

20. Crean RM, Pudney CR, Cole DK, van der Kamp MW. Reliable In Silico Ranking of Engineered Therapeutic TCR Binding Affinities with MMPB/GBSA. J Chem Inf Model. 2022;62(3):577–90.

21. Peacock T, Chain B. Information-Driven Docking for TCR-pMHC Complex Prediction. Front Immunol. 2021 Sep;12:686127.

22. Zhu Y, Huang C, Su M, Ge Z, Gao L, Shi Y, et al. Characterization of amino acid residues of T-cell receptors interacting with HLA-A*02-restricted antigen peptides. Ann Transl Med. 2021 Mar;9(6):495.

23. Wu D, Efimov GA, Bogolyubova AV, Pierce BG, Mariuzza RA. Structural insights into protection against a SARS-CoV-2 spike variant by T cell receptor (TCR) diversity. J Biol Chem. 2023;103035.

24. Szeto C, Nguyen AT, Lobos CA, Chatzileontiadou DSM, Jayasinghe D, Grant EJ, et al. Molecular Basis of a Dominant SARS-CoV-2 Spike-Derived Epitope Presented by HLA-A*02:01 Recognised by a Public TCR. Cells [Internet]. 2021 Mar;10(10). Available from: http://dx.doi.org/10.3390/cells10102646

25. Chaurasia P, Nguyen THO, Rowntree LC, Juno JA, Wheatley AK, Kent SJ, et al. Structural basis of biased T cell receptor recognition of an immunodominant HLA-A2 epitope of the SARS-CoV-2 spike protein. J Biol Chem. 2021 Jan;297(3):101065.

26. Wu D, Kolesnikov A, Yin R, Guest JD, Gowthaman R, Shmelev A, et al. Structural assessment of HLA-A2-restricted SARS-CoV-2 spike epitopes recognized by public and private T-cell receptors. Nat Commun. 2022 Oct;13(1):19.

27. Davis MM, Bjorkman PJ. T-cell antigen receptor genes and T-cell recognition. Nature. 1988 Apr;334(6181):395–402.

28. Krogsgaard M, Davis MM. How T cells “see” antigen. Nat Immunol. 2005 Mar;6(3):239–45.

29. Springer I, Besser H, Tickotsky-Moskovitz N, Dvorkin S, Louzoun Y. Prediction of Specific TCR-Peptide Binding From Large Dictionaries of TCR-Peptide Pairs. Front Immunol. 2020;11:1803.

30. Shugay M, Bagaev DV, Zvyagin IV, Vroomans RM, Crawford JC, Dolton G, et al. VDJdb: a curated database of T-cell receptor sequences with known antigen specificity. Nucleic Acids Res. 2018 Apr;46(D1):D419–27.

31. Tynan FE, Reid HH, Kjer-Nielsen L, Miles JJ, Wilce MCJ, Kostenko L, et al. A T cell receptor flattens a bulged antigenic peptide presented by a major histocompatibility complex class I molecule. Nat Immunol. 2007 Mar;8(3):268–76.

32. Bagaev DV, Vroomans RMA, Samir J, Stervbo U, Rius C, Dolton G, et al. VDJdb in 2019: database extension, new analysis infrastructure and a T-cell receptor motif compendium. Nucleic Acids Res. 2020 Aug;48(D1):D1057–62.

33. Borrman T, Cimons J, Cosiano M, Purcaro M, Pierce BG, Baker BM, et al. ATLAS: A database linking binding affinities with structures for wild-type and mutant TCR-pMHC complexes. Proteins. 2017 May;85(5):908–16.

34. Atchley WR, Zhao J, Fernandes AD, Drüke T. Solving the protein sequence metric problem. Proc Natl Acad Sci U S A. 2005 Mar;102(18):6395–400.

35. Leaver-Fay A, Tyka M, Lewis SM, Lange OF, Thompson J, Jacak R, et al. ROSETTA3: an object-oriented software suite for the simulation and design of macromolecules. Methods Enzymol. 2011;487:545–74.

36. Martin AJM, Vidotto M, Boscariol F, Di Domenico T, Walsh I, Tosatto SCE. RING: networking interacting residues, evolutionary information and energetics in protein structures. Bioinformatics. 2011;27(14):2003–5.

37. Osorio D, Rondón-Villarreal P, Torres R. Peptides: A package for data mining of antimicrobial peptides. R J. 2015;7(1):4.

38. R Core Team. R: A language and environment for statistical computing. R Foundation for Statistical Computing. Available from: https://www.R-project.org/

39. Lindahl, Abraham, Hess, Spoel V der. GROMACS 2020 Manual [Internet]. 2020. Available from: https://zenodo.org/record/3562512

40. McLachlan AD. Rapid comparison of protein structures. Acta Crystallogr A. 1982 Jan;38(6):871–3.

